# Sympathetic innervation of interscapular brown adipose tissue is not a predominant mediator of oxytocin-elicited reductions of body weight and adiposity in male diet-induced obese mice

**DOI:** 10.1101/2024.05.29.596425

**Authors:** Melise M. Edwards, Ha K. Nguyen, Andrew D. Dodson, Adam J. Herbertson, Tami Wolden-Hanson, Tomasz Wietecha, Mackenzie K. Honeycutt, Jared D. Slattery, Kevin D. O’Brien, James L. Graham, Peter J. Havel, Thomas O. Mundinger, Carl Sikkema, Elaine R. Peskind, Vitaly Ryu, Gerald J. Taborsky, James E. Blevins

## Abstract

Previous studies indicate that CNS administration of oxytocin (OT) reduces body weight in high fat diet-induced obese (DIO) rodents by reducing food intake and increasing energy expenditure (EE). We recently demonstrated that hindbrain (fourth ventricular [4V]) administration of OT elicits weight loss and elevates interscapular brown adipose tissue temperature (T_IBAT_, a surrogate measure of increased EE) in DIO mice. What remains unclear is whether OT-elicited weight loss requires increased sympathetic nervous system (SNS) outflow to IBAT. We hypothesized that OT-induced stimulation of SNS outflow to IBAT contributes to its ability to activate BAT and elicit weight loss in DIO mice. To test this hypothesis, we determined the effect of disrupting SNS activation of IBAT on the ability of 4V OT administration to increase T_IBAT_ and elicit weight loss in DIO mice. We first determined whether bilateral surgical SNS denervation to IBAT was successful as noted by ≥ 60% reduction in IBAT norepinephrine (NE) content in DIO mice. NE content was selectively reduced in IBAT at 1-, 6- and 7-weeks post-denervation by 95.9±2.0, 77.4±12.7 and 93.6±4.6% (*P*<0.05), respectively and was unchanged in inguinal white adipose tissue, pancreas or liver. We subsequently measured the effects of acute 4V OT (1, 5 µg ≈ 0.99, 4.96 nmol) on T_IBAT_ in DIO mice following sham or bilateral surgical SNS denervation to IBAT. We found that the high dose of 4V OT (5 µg ≈ 4.96 nmol) elevated T_IBAT_ similarly in sham mice as in denervated mice. We subsequently measured the effects of chronic 4V OT (16 nmol/day over 29 days) or vehicle infusions on body weight, adiposity and food intake in DIO mice following sham or bilateral surgical denervation of IBAT. Chronic 4V OT reduced body weight by 5.7±2.23% and 6.6±1.4% in sham and denervated mice (*P*<0.05), respectively, and this effect was similar between groups (*P*=NS). OT produced corresponding reductions in whole body fat mass (*P*<0.05). Together, these findings support the hypothesis that sympathetic innervation of IBAT is not necessary for OT-elicited increases in BAT thermogenesis and reductions of body weight and adiposity in male DIO mice.

## Introduction

While the hypothalamic neuropeptide, oxytocin (OT), has a more clearly established role in the control of reproductive behavior [1] and prosocial behavior [2; 3], it has also been recognized as having an important role in the control of food intake and body weight [4; 5; 6; 7]. Although OT reduces body weight in part by decreasing energy intake, pair-feeding studies indicate that OT-elicited reductions of weight gain or weight loss cannot be fully explained by its ability to decrease energy intake [8; 9; 10]. In these studies, pair-feeding appears to account for approximately 50% of the reduction of weight gain and/or weight loss observed with OT treatment [8; 9; 10]

Indeed, recent studies have shown that OT increases energy expenditure (EE) [11; 12; 13; 14]. While brown adipose tissue thermogenesis (BAT) is important in the control of EE [for review see [15; 16]], it is unclear whether OT’s effects on EE result primarily from 1) activation of non-shivering BAT thermogenesis, 2) spontaneous physical activity-induced thermogenesis [17] 3) shivering and non-shivering thermogenesis in skeletal muscle [18], or 4) endocrine factors (e.g. thyroid hormone, fibroblast growth factor-21, irisin (for review see [19; 20])). We have found that acute third (3V) and fourth ventricular (4V) injections of OT increase interscapular BAT temperature (T_IBAT_), a functional readout of BAT thermogenesis in mice and rats [21; 22]. Furthermore, the effects of chronic CNS administration of OT on T_IBAT_ coincide with OT-elicited weight loss in diet-induced obese (DIO) rats [21]. Sutton and colleagues demonstrated that use of designer drugs (DREADDs) technology to chemogenetically activate hypothalamic paraventricular nucleus (PVN) OT neurons increases both EE and tends to increase subcutaneous BAT temperature (*P*=0.13) in *Oxytocin-Ires-Cre* mice [23]. Furthermore, Yuan et al recently reported that peripheral administration of OT promotes BAT differentiation *in vitro* and the expression of genes involved in thermogenesis in IBAT in high fat diet-fed mice [24]. On the other hand, reduced OT signaling is associated with obesity [14; 25; 26; 27], reductions of EE [13; 14; 27; 28] and deficits in BAT thermogenesis [28; 29; 30; 31] in mice. Collectively, these findings support a role for increased BAT thermogenesis in OT-elicited weight loss in mice. What remains unclear is whether OT-elicited weight loss requires increased sympathetic nervous system (SNS) outflow to IBAT and whether this effect involves hindbrain OT receptors (OTRs). Here, we sought to clarify the role of SNS outflow to IBAT in mediating the effects of hindbrain OTR stimulation on both weight loss and BAT thermogenesis.

We hypothesized that OT-induced stimulation of SNS outflow to IBAT contributes to its ability to stimulate non-shivering BAT thermogenesis and elicit weight loss in DIO mice. We first confirmed the success of the IBAT denervation procedure by measuring IBAT norepinephrine (NE) content at 1-week post-denervation in lean mice. We subsequently determined if these effects can be translated to DIO mice at 1, 6 and 7-weeks post-sham/denervation. To assess the role of SNS innervation of BAT in contributing to OT-elicited increases in non-shivering thermogenesis in IBAT (as surrogate measure of energy expenditure), we examined the effects of acute 4V OT (1, 5 μg) on T_IBAT_ in DIO mice following bilateral surgical SNS denervation to IBAT. We next determined if SNS innervation to IBAT contributes to OT-induced weight loss by measuring the effect of bilateral surgical or sham denervation of IBAT on the ability of chronic 4V OT (16 nmol/day over 29 days) or vehicle administration to reduce body weight and adiposity. We subsequently determined if these effects were associated with a reduction of adipocyte size and energy intake.

## Methods

### Animals

Adult male C57BL/6J mice were initially obtained from Jackson Laboratory [strain # 000664 (lean) or 380050 (DIO); Bar Harbor, ME)] after having been maintained on a chow diet (∼16 weeks; 25-35 grams upon arrival) or the high fat diet (HFD) for 4 months starting at 6 weeks of age (∼22 weeks; 40-50 grams upon arrival). The chow diet provided 16% kcal from fat (∼0.62% sucrose) (5LG4; LabDiet®, St. Louis, MO). The HFD provided 60% kcal from fat (∼ 6.8% kcal from sucrose and 8.9% of the diet from sucrose) (Research Diets, Inc., D12492i, New Brunswick, NJ). All animals were housed individually in Plexiglas cages in a temperature-controlled room (22±2°C) under a 12:12-h light-dark cycle. All mice were maintained on a 6 a.m./6 p.m. light cycle. Mice had *ad libitum* access to water and HFD. The research protocols were approved both by the Institutional Animal Care and Use Committee of the Veterans Affairs Puget Sound Health Care System (VAPSHCS) and the University of Washington in accordance with NIH Guidelines for the Care and Use of Animals.

### Drug Preparation

Fresh solutions of OT acetate salt were prepared the day of each experiment (**Study 6**). Fresh solutions of OT acetate salt (Bachem Americas, Inc., Torrance, CA) were solubilized in sterile water, loaded into Alzet® minipumps (model 2004; DURECT Corporation, Cupertino, CA) and subsequently primed in sterile 0.9% saline at 37° C for approximately 40 hours prior to minipump implantation based on manufacturer’s recommended instructions (**Study 4-5**). The beta-3 receptor (β-3R) agonist, CL 316243 (Tocris/Bio-Techne Corporation, Minneapolis, MN), was solubilized in sterile water each day of each experiment (**Study 3**).

### SNS denervation procedure

A dissecting microscope (Leica M60/M80; Leica Microsystems, Buffalo Grove, IL) was used throughout the procedure. A 1” midline incision was made in the skin dorsally at the level of the thorax and continued rostrally to the base of the skull. Connective tissue was blunt dissected away from the adipose tissue with care to avoid cutting the large thoracodorsal artery that is located medially to both pads. Both left and right fat pads were separated from the mid line. Each fat pad was lifted up and the intercostal nerve bundles were located below. Once the nerves were located, a sharp point forceps was used to pull the nerve bundles straight up while using a 45 degree scissors to cut and remove 3-5 mm of nerves. The interscapular incision was closed with 4-0 non-absorbable monofilament Ethilon (nylon) sutures or with standard metal wound clips. Nerves were similarly identified but not cut for sham operated animals. Mice were treated pre-operatively with the analgesic ketoprofen (2 mg/kg; Fort Dodge Animal Health) prior to the completion of the denervation or sham procedure. This procedure was combined with transponder implantations for studies that involved IBAT temperature measurements in response to acute 4V injections (**Study 4**). Animals were allowed to recover for approximately 5-7 days prior to implantation of 4V cannulas. Note that surgical denervation was used in place of chemical denervation because chemical denervation can result in SNS terminal recovery within IBAT [32; 33] within only 10 days post-chemical denervation [32].

### 4V cannulations for acute injections

Animals were implanted with a cannula (P1 Technologies, Roanoke, VA) that was directed towards the 4V as previously described [22]. Briefly, mice under isoflurane anesthesia were placed in a stereotaxic apparatus with the incisor bar positioned 4.5 mm below the interaural line. A 26-gauge cannula (P1 Technologies) was stereotaxically positioned into the 4V (5.9 mm caudal to bregma; 0.4 mm lateral to the midline, and 2.7 mm ventral to the skull surface) and secured to the surface of the skull with dental cement and stainless-steel screws.

### 4V cannulations for chronic infusions

Mice were implanted with a cannula within the 4V with a side port that was connected to an osmotic minipump (model 2004, DURECT Corporation) as previously described [22]. Mice under isoflurane anesthesia were placed in a stereotaxic apparatus with the incisor bar positioned 4.5 mm below the interaural line. A 30-gauge cannula (P1 Technologies) was stereotaxically positioned into the 4V (5.9 mm caudal to bregma; 0.4 mm lateral to the midline, and 3.7 mm ventral to the skull surface) and secured to the surface of the skull with dental cement and stainless steel screws. A 1.2” piece of plastic Tygon^TM^ Microbore Tubing (0.020” x 0.060“OD; Cole-Parmer) was tunneled subcutaneously along the midline of the back and connected to the 21-gauge sidearm osmotic minipump-cannula assembly. A stainless steel 22-gauge pin plug (Instech Laboratories, Inc.) was temporarily inserted at the end of the tubing during a two week postoperative recovery period, after which it was replaced by an osmotic minipump (DURECT Corporation) containing saline or OT. Mice were treated with the analgesic ketoprofen (5 mg/kg; Fort Dodge Animal Health) and the antibiotic enrofloxacin (5 mg/kg; Bayer Healthcare LLC., Animal Health Division Shawnee Mission, KS) at the completion of the 4V cannulations and were allowed to recover at least 10 days prior to implantation of osmotic minipumps.

### Implantation of temperature transponders underneath IBAT

Animals were anesthetized with isoflurane and had the dorsal surface along the upper midline of the back shaved. The area was subsequently scrubbed with 70% ethanol followed by betadine swabs to sterilize/clean the area before a one-inch incision was made at the midline of the interscapular area. The temperature transponder (14 mm long/2 mm wide) (HTEC IPTT-300; BIO MEDIC DATA SYSTEMS, INC, Seaford, DE) was implanted underneath both IBAT pads as previously described [22] and secured in place by suturing it to the brown fat pad with sterile silk suture. HTEC IPTT-300 transponders were used in place of IPTT-300 transponders to enhance accuracy in our measurements. The IPTT-300 transponders are specified to be accurate to ± 0.4°C between 35°C and 39°C (± 1.0°C between 32°C and 42°C) while the HTEC IPTT-300 transponders are accurate to ± 0.2°C between 32°C and 42°C (personal communication with Geoff Hunt from BIO MEDIC DATA SYSTEMS). The interscapular incision was closed with Nylon sutures (5-0), which were removed in awake animals 10-14 days after surgery.

### Acute IP injections and measurements of T_IBAT_

OT (or saline vehicle; 1 μL injection volume) was administered immediately prior to the start of the dark cycle following 4 hours of food deprivation. Animals remained without access to food for an additional 4 h during the T_IBAT_ measurements to prevent the confounding effects of diet-induced thermogenesis on T_IBAT_. A handheld reader (DAS-8007-IUS Reader System; BIO MEDIC DATA SYSTEMS, INC) was used to collect measurements of T_IBAT_. The 4V injections were administered at 1 μL/min using an injection pump (Harvard Apparatus Pump II Elite; Harvard Apparatus, Holliston, MA) via a 33-gauge injector (P1 Technologies) connected by polyethylene 20 tubing to a 10-μL Hamilton syringe. Mice underwent all treatments (unless otherwise noted) in a randomized order separated by at least 48 hours between treatments.

### Body Composition

Determinations of lean body mass and fat mass were made on un-anesthetized mice by quantitative magnetic resonance using an EchoMRI 4-in-1-700^TM^ instrument (Echo Medical Systems, Houston, TX) at the VAPSHCS Rodent Metabolic Phenotyping Core. Measurements were taken prior to 4V cannulations and minipump implantations as well as at the end of the infusion period.

## Tissue collection for NE content measurements

Mice were euthanized by rapid conscious decapitation at 1-week (**Studies 1-3**), 6-weeks (**Study 2**), 7-weeks (**Study 2, Study 6**), 9-weeks (**Study 4**) or 10-11 weeks (**Study 5**) post-sham or denervation procedure. Trunk blood and tissues (IBAT, EWAT, IWAT, liver or pancreas) were collected from 4-h fasted mice. Tissue was rapidly removed, wrapped in foil and frozen in liquid N2. Samples were stored frozen at −80°C until analysis. Note that anesthesia was not used when collecting tissue for NE content as it can cause the release of NE from SNS terminals within the tissue [33].Thus, animals were euthanized by rapid conscious decapitation.

### Norepinephrine (NE) content measurements (biochemical confirmation of IBAT denervation procedure)

NE content was measured in IBAT, EWAT, IWAT, liver and/or pancreas using previously established techniques [34]. Successful denervation was noted by ≥ 60% reduction in IBAT NE content as previously noted [35]. Experimental animals that did not meet this criterion were excluded from the data analysis.

## Study Protocols

### Study 1A: Determine the success of the surgical denervation procedure at 1-week post-sham or denervation in lean mice by measuring NE content

Mice underwent sham or bilateral surgical SNS denervation procedures and, to prevent the confounder of anesthesia on NE content, animals were euthanized by rapid conscious decapitation at 1-week post-sham or denervation procedure.

### Study 1B: Determine the success of the surgical denervation procedure at 10-12 weeks post-sham or denervation in lean mice by measuring thermogenic gene expression

Mice underwent sham or bilateral surgical SNS denervation procedures and were euthanized by rapid conscious decapitation at 1-week post-sham or denervation procedure.

### Study 2: Determine the success of the surgical denervation procedure at 1-, 6- and 7-weeks post-sham or denervation in DIO mice by measuring NE content

Mice were fed *ad libitum* and maintained on HFD for approximately 4.25 months prior to underdoing sham or SNS denervation procedures. In addition to weekly body weight measurements, body composition measurements were obtained at baseline and at 6 and 7-weeks post-denervation/sham procedures. Mice were euthanized by rapid conscious decapitation 1-, 6- and 7-weeks post-sham or denervation procedure.

### Study 3: Determine if SNS innervation of IBAT is reduced in age-matched obese mice relative to lean mice by measuring NE content

Chow-fed and HFD-fed mice were age-matched (∼22 weeks; 30-35 grams or 40-50 grams upon arrival). Age-matched chow-fed and DIO mice were fed *ad libitum* and maintained on chow or HFD for 4.5 months and matched for both body weight and adiposity prior to undergoing sham (N=5/diet) or SNS denervation (N=5/diet) procedures. Mice were euthanized by rapid conscious decapitation at 1-week post-sham or denervation procedure.

### Study 4: Determine if surgical denervation of IBAT compromises the ability of the beta-3 receptor agonist, CL 316243, to increase T_IBAT_ in DIO mice

Mice were fed *ad libitum* and maintained on HFD for 4.5 months prior to underdoing sham or SNS denervation procedures and implantation of temperature transponders underneath the left IBAT depot. Mice were allowed to recover for at least 1 week during which time they were adapted to a daily 4-h fast, handling and mock injections. On an experimental day, 4-h fasted mice received CL 316243 (0.1 or 1 mg/kg; IP) or vehicle (sterile water; 1.5 mL/kg injection volume) during the early part of the light cycle in order to maximize the effects of CL 316243 during a time when circulating NE levels [36] and IBAT catecholamine levels are lower [37]. Injections were completed in a crossover design at 7-day intervals such that each animal served as its own control (approximately 4-6 weeks post-sham or denervation procedures). T_IBAT_ was measured at baseline (−2 h; 8:00 a.m.), immediately prior to IP injections (0 h; 9:45-10:00 a.m.), and at 0.25, 0.5, 0.75, 1, 1.25, 1.5, 2, 3, 4, and 24-h post-injection (9:45-10:00 a.m.). Food intake and body weight were measured daily. This dose range was based on doses of CL 316243 found to be effective at reducing food intake and weight gain in rats [22; 38] (IP) and stimulating IBAT temperature in rats [22; 39] (IV, IP) and mice (IV) [40]. Animals were euthanized by rapid conscious decapitation at 9-weeks post-sham or denervation procedure. Similar studies were also performed in lean mice (data not shown).

Likewise, we also examined the effects of systemic administration of the transient receptor potential channel melastatin family member 8 (TRPM8) agonist, icilin, which, may activate IBAT through a direct [41] or indirect mechanism [42] (data not shown). In addition, the effects of the sympathomimetic, tyramine, was also examined on T_IBAT_ in DIO mice with intact or impaired SNS innervation of IBAT (**Supplemental Study 1**).

### Study 5: Determine the extent to which 4V OT-induced activation of sympathetic outflow to IBAT contributes to its ability to increase T_IBAT_ in DIO mice

Mice were fed *ad libitum* and maintained on HFD for 4.25 months prior to undergoing sham or SNS denervation procedures and implantation of temperature transponders underneath the left IBAT depot. Mice were subsequently implanted 1 week later with 4V cannulas. Mice were allowed to recover for at least 2 weeks during which time they were adapted to a daily 4-h fast, handling and mock injections. On an experimental day, 4-h fasted mice received OT (1 or 5 μg/μl) or vehicle during the early part of the light cycle in order to maximize the effects of OT [14; 22] during a time when circulating NE levels [36] and IBAT catecholamine levels are lower [37]. Injections were completed in a crossover design at approximately 48-h intervals such that each animal served as its own control (approximately 7-8 weeks post-sham or denervation procedures). T_IBAT_ was measured at baseline (−2 h; 9:00 a.m.), immediately prior to 4V injections (0 h; 9:45-10:00 a.m.), and at 0.25, 0.5, 0.75, 1, 1.25, 1.5, 2, 3, 4, and 24-h post-injection (10:00 a.m.). Food intake and body weight were measured daily. This dose range was based on doses of 4V OT found to be effective at stimulating T_IBAT_ in DIO mice in previous studies [22]. Mice were euthanized by rapid conscious decapitation at 1-week post-sham or denervation procedure.

Similar studies were also performed in lean mice (**Supplemental Study 2**). In addition, we examined the effectiveness of a systemic dose to recapitulate the effects of a 4V dose to increase T_IBAT_ in lean mice (**Supplemental Study 3**). We next determined if the melanocortin 3/4 receptor (MC3/4R) agonist, melanotan II (MTII), which, along with the endogenous MC3/4R agonist, alpha-MSH, activate PVN OT neurons [43; 44] and act, in part, through OT signaling [43; 45], could recapitulate the effects of systemic OT on T_IBAT_ in lean mice with intact or impaired SNS innervation of IBAT (data not shown).

### Study 6A: Determine the extent to which OT-induced activation of sympathetic outflow to IBAT contributes to its ability to elicit weight loss in DIO mice

Mice were fed *ad libitum* and maintained on HFD for 4.25-4.5 months prior to prior to undergoing sham or SNS denervation procedures and implantation of temperature transponders underneath the left IBAT depot. Mice subsequently received 4V cannulas and minipumps to infuse vehicle or OT (16 nmol/day) over 28 days as previously described [22]. This dose was selected based on a dose of 4V OT found to be effective at reducing body weight in DIO mice [22]. Daily food intake and body weight were also tracked for 28 days. Animals were euthanized by rapid conscious decapitation at 7 weeks post-sham or denervation procedure.

### Study 6B: Determine the extent to which OT-induced activation of sympathetic outflow to IBAT impacts thermogenic gene expression in IBAT, IWAT and EWAT in DIO mice

Mice from **Study 6A** were used for these studies. All mice received injections of 4V vehicle or OT and were euthanized by rapid conscious decapitation at 2-h post-injection.

### Study 7: Determine the extent to which systemic (subcutaneous) infusion of a centrally effective dose of OT (16 nmol/day) elicits weight loss in DIO mice

Mice were fed *ad libitum* and maintained on HFD for 4-4.25 months prior to prior to being implanted with a temperature transponder underneath the left IBAT depot. Mice were subsequently maintained on a daily 4-h fast and received minipumps to infuse vehicle or OT (16 nmol/day) over 28 days [22]. This dose was selected based on a dose of 4V OT found to be effective at reducing body weight in DIO mice [22]. Daily food intake and body weight were also tracked for 28 days. Mice were euthanized with intraperitoneal injections of ketamine cocktail [ketamine hydrochloride (390 mg/kg), xylazine (26.4mg/kg) in an injection volume up to 1 mL/mouse] prior to collection of blood (cardiac stick) and tissues (IBAT, EWAT, IWAT, gastrocnemius and brain). Tissue was otherwise treated identically to Studies 1-4 or processed for subsequent measurements of adipocyte size and UCP-1 (see below).

### Blood collection

Blood samples [trunk blood (**Study 3-6**) or cardiac stick (**Study 7**)] were collected from 4-h fasted mice within a 2-h window towards the beginning of the light cycle (10:00 a.m.-12:00 p.m.) as previously described in DIO CD^®^ IGS and Long-Evans rats and C57BL/6J mice [21; 22; 46]. Treatment groups were counterbalanced at time of euthanasia to avoid time of day bias. Blood samples [up to 1 mL] were collected by trunk blood or cardiac stick in chilled K2 EDTA Microtainer Tubes (Becton-Dickinson, Franklin Lakes, NJ). Whole blood was centrifuged at 6,000 rpm for 1.5-min at 4°C; plasma was removed, aliquoted and stored at −80°C for subsequent analysis.

### Plasma hormone measurements

Plasma leptin and insulin were measured using electrochemiluminescence detection [Meso Scale Discovery (MSD^®^), Rockville, MD] using established procedures [21; 47]. Intra-assay coefficient of variation (CV) for leptin was 4.2% and 1.9% for insulin. The range of detectability for the leptin assay is 0.137-100 ng/mL and 0.069-50 ng/mL for insulin. Plasma fibroblast growth factor-21 (FGF-21) (R&D Systems, Minneapolis, MN) and irisin (AdipoGen, San Diego, CA) levels were determined by ELISA. The intra-assay CV for FGF-21 and irisin were 2.3% and 9.3%, respectively; the ranges of detectability were 31.3-2000 pg/mL (FGF-21) and 0.078-5 μg/mL (irisin). Plasma adiponectin was determined by ELISA [Millipore Sigma (Burlington, MA)]. Intra-assay CV for adiponectin was 4.2%. The range of detectability for the adiponectin assay is 2.8-178 ng/mL. The data were normalized to historical values using a pooled plasma quality control sample that was assayed in each plate.

### Blood glucose and lipid measurements

Blood was collected for glucose measurements by tail vein nick in 4-h fasted rats and measured with a glucometer using the AlphaTRAK 2 blood glucose monitoring system (Abbott Laboratories, Abbott Park, IL) [21; 48]. Total cholesterol (TC) [Fisher Diagnostics (Middletown, VA)], free fatty acids (FFAs) [Wako Chemicals USA, Inc., Richmond, VA)] and free glycerol (FG) (Millipore Sigma) were measured using an enzymatic-based kits. Intra-assay CVs for TC, FFAs and FG were 1.5, 2.9, and 2.2%, respectively. These assay procedures have been validated for rodents [49].

### Adipose tissue processing for adipocyte size

Adipose tissue depots were collected at the end of the infusion period from a subset of DIO mice from **Study 6** (IBAT, IWAT, EWAT) and **Study 7** (EWAT and IWAT). IBAT, IWAT, and EWAT depots were dissected and placed in 4% paraformaldehyde-PBS for 24 h and then placed in 70% ethanol (EtOH) prior to paraffin embedding. Sections (5 μm) sampled were obtained using a rotary microtome, slide-mounted using a floatation water bath (37°C), and baked for 30 min at 60°C to give approximately 15-16 slides/fat depot with two sections/slide.

### Adipocyte size analysis

Adipocyte size analysis was performed on deparaffinized and digitized IWAT and EWAT sections. The average cell area from two randomized photomicrographs was determined using the built-in particle counting method of ImageJ software (National Institutes of Health, Bethesda, MD). Fixed (4% PFA), paraffin-embedded adipose tissue was sectioned. Slides were visualized using bright field on an Olympus BX51 microscope (Olympus Corporation of the Americas; Center Valley, PA) and photographed using a Canon EOS 5D SR DSLR (Canon U.S.A., Inc., Melville, NY) camera at 100X magnification. Values for each tissue within a treatment were averaged to obtain the mean of the treatment group.

### Tissue collection for NE content measurements

Mice were euthanized by rapid conscious decapitation at 1-week (**Studies 1-3**), 6-weeks (**Study 2**), 7-weeks (**Study 2, Study 6**), 9-weeks (**Study 4**) or 10-11 weeks (**Study 5**) post-sham or denervation procedure. Trunk blood and tissues (IBAT, EWAT, IWAT, liver or pancreas) were collected from 4-h fasted mice. Tissue was rapidly removed, wrapped in foil and frozen in liquid N2. Samples were stored frozen at −80°C until analysis. Note that anesthesia was not used when collecting tissue for NE content as it can cause the release of NE from SNS terminals within the tissue [33].

### Tissue collection for quantitative real-time PCR (qPCR)

Tissue (IBAT, IWAT and EWAT) was collected from a subset of 4-h fasted mice in **Study 4-7**. In addition, mice from **Study 7** were euthanized with an overdose of ketamine cocktail prior to tissue collection. IBAT, IWAT, EWAT, gastrocnemius and brains were collected within a 2-h window towards the start of the light cycle (10:00 a.m.-12:00 p.m.) as previously described in DIO CD^®^ IGS/Long-Evans rats and C57BL/6J mice [21; 22; 46]. Tissue was rapidly removed, wrapped in foil and frozen in liquid N2. Samples were stored frozen at −80°C until analysis.

### qPCR

RNA extracted from samples of IBAT and IWAT (**Studies 4-7**) were analyzed using the RNeasy Lipid Mini Kit (Qiagen Sciences Inc, Germantown, MD) followed by reverse transcription into cDNA using a high-capacity cDNA archive kit (Applied Biosystems, Foster City, CA). Quantitative analysis for relative levels of mRNA in the RNA extracts was measured in duplicate by qPCR on an Applied Biosystems 7500 Real-Time PCR system (Thermo Fisher Scientific, Waltham, MA) and normalized to the cycle threshold value of Nono mRNA in each sample. The TaqMan® probes used in the study were Thermo Fisher Scientific Gene Expression Assay probes. The probe for mouse Nono (Mm00834875_g1), UCP-1 (catalog no. Mm01244861_m1), UCP-2 (catalog no. Mm00627599_m1), UCP-3 (catalog no. Mm01163394_m1), beta-1 adrenergic receptor (Adrb1; catalog no. Mm00431701_s1), beta-2 adrenergic receptor (Adrb2; catalog no. Mm02524224_s1), beta-3 adrenergic receptor (Adrb3; catalog no. Mm02601819_g1), alpha-2 adrenergic receptor (Adra2a; catalog no. Mm07295458_s1),type 2 deiodinase (D2) (Dio2; catalog no. Mm00515664_m1), cytochrome c oxidase subunit 8b (Cox8b; catalog no. Mm00432648_m1), G-protein coupled receptor 120 (Gpr120; catalog no. Mm00725193_m1), bone morphogenetic protein 8b (bmp8b; catalog no. Mm00432115_g1), cell death-inducing DNA fragmentation factor alpha-like effector A (Cidea; catalog no. Mm00432554_m1), peroxisome proliferator-activated receptor gamma coactivator 1 alpha (Ppargc1a; catalog no. Mm01208835_m1) were acquired from Thermo Fisher Scientific. Relative amounts of target mRNA were determined using the Comparative C_T_ or 2-^ΔΔC^_T_ method [50] following adjustment for the housekeeping gene.

### Transponder placement

All temperature transponders were confirmed to have remained underneath the IBAT depot at the conclusion of the study.

### Statistical Analyses

All results are expressed as means ± SE. Comparisons between multiple groups involving between subjects design were made using one-way ANOVA as appropriate, followed by a post-hoc Fisher’s least significant difference test. Comparisons involving within-subjects designs were made using a one-way repeated-measures ANOVA followed by a post-hoc Fisher’s least significant difference test. Analyses were performed using the statistical program SYSTAT (Systat Software, Point Richmond, CA). Differences were considered significant at *P*<0.05, 2-tailed.

## Results

### Study 1A: Determine the success of the surgical denervation procedure at 1-week post-sham or denervation in lean mice

The goal of this study was to verify the success of the SNS denervation of IBAT procedure in lean mice by confirming a reduction of NE content that was specific to IBAT relative to other tissues (IWAT and liver). By design, mice were lean as determined by body weight (25.4±0.3 g).

IBAT NE content was reduced by 99.6±0.2% at 1-week post-denervation [(F(1,8) = 12.358, *P*=0.008)] (**Figure 1**) relative to IBAT from sham operated mice. In contrast, NE content was unchanged in IWAT or liver in denervated mice relative to sham mice (*P*=NS). We also repeated this study in a separate group of lean mice and found a similar reduction of IBAT NE content (99.2±0.4%) at 1-week post-denervation relative to sham operated mice [(F(1,8) = 73.438, *P*=0.000)].

**Figure 1.**
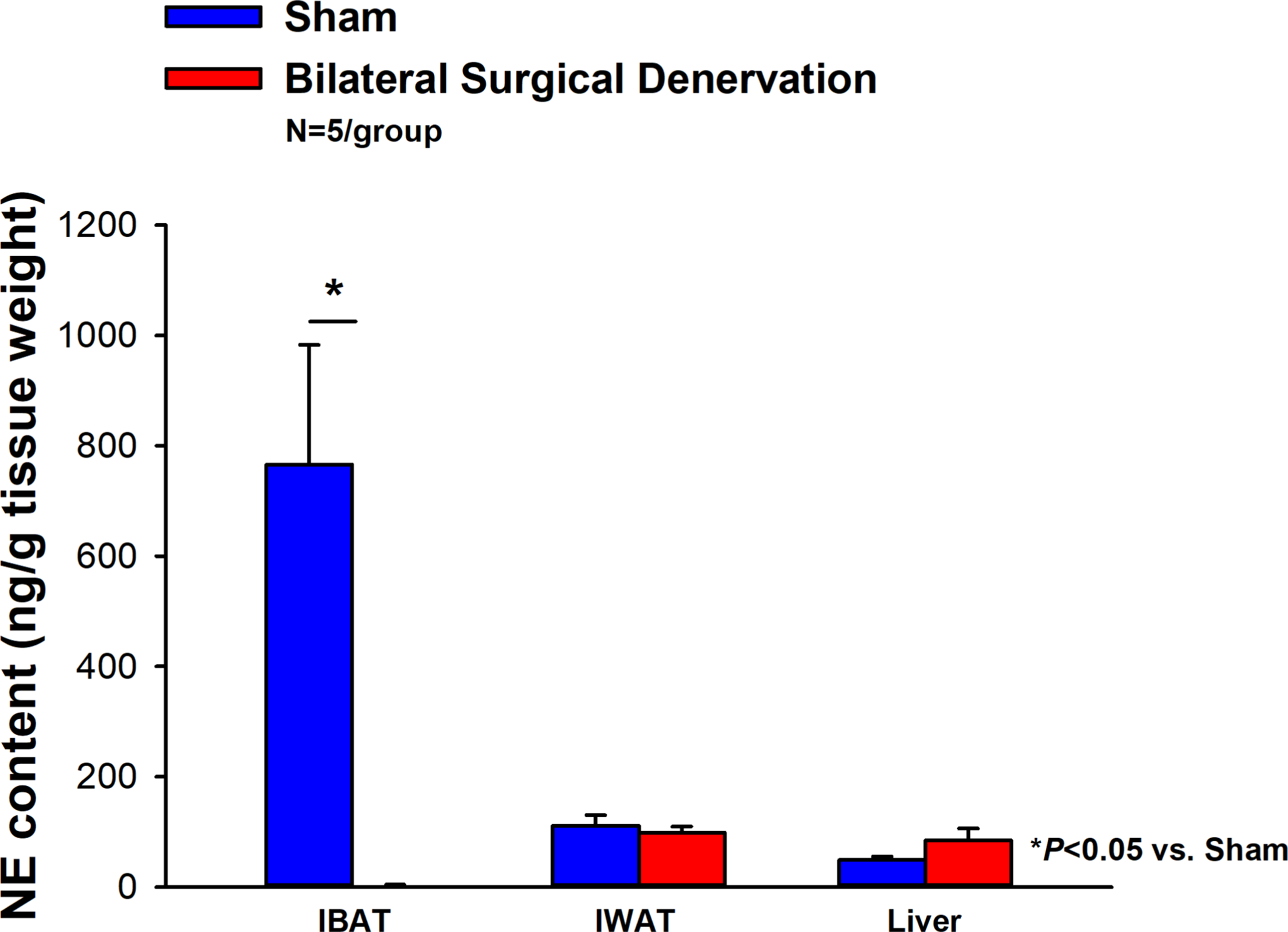
Effect of IBAT surgical denervation procedure on IBAT NE content at 1-week post-sham or IBAT denervation in lean mice. Mice were maintained on chow for approximately 2.5 months prior to undergoing a sham (N=5/group) or bilateral surgical IBAT denervation (N=5/group) procedures and were euthanized at 1-week post-sham or denervation procedure. NE content was measured from IBAT, IWAT and liver. Data are expressed as mean ± SEM. \**P*<0.05, †0.05<*P*<0.1 denervation vs sham.

### Study 1B: Verify the success of the surgical denervation procedure at 10-12 weeks after IBAT surgical denervation or sham denervation in lean mice by measuring thermogenic gene expression

The goal of this study was to determine the 1) verify the success of the SNS denervation of IBAT procedure in lean mice using a separate biochemical marker (IBAT thermogenic gene expression) and 2) confirm that these changes would persist for extended periods of time out to 10-12 weeks.

IBAT NE content was reduced in denervated mice by 96.1±3.3% at 10-12 weeks after IBAT surgical denervation relative to sham-operated control mice [(F(1,2) = 87.393, *P*=0.011). Only a subset of samples (N=4 out of 20) were able to be screened for NE content but all tissue samples were included in the IBAT (N=20 out of 20), IWAT (N=9 out of 20) and EWAT (N=9 out of 20) gene expression analysis.

#### IBAT

There was a reduction of 9 out of the 13 measured mRNA levels of genes from IBAT of denervated mice relative to IBAT from sham operated mice (*P*<0.05; Table 1A).There were significant reductions of IBAT UCP-1 [(F(1,7) = 38.1, *P*<0.001)], Dio2 [(F(1,7) = 12.669, *P*=0.009)], Gpr120 [(F(1,7) = 65.965, *P*<0.001)], Adrb3 [(F(1,7) = 65.916, *P*<0.001)], Adrb1 [(F(1,7) = 8.015, *P*=0.025)], Acox1 [(F(1,7) = 58.261, *P*<0.001)], bmp8b [(F(1,7) = 19.636, *P*=0.003)], cox8b [(F(1,6) = 16.403, *P*=0.007)] and UCP-3 mRNA expression [(F(1,7) = 57.665, *P*<0.001)]. There were no significant differences in IBAT CIDEA, PPARGC1, PPARA and PRDM16 mRNA expression between sham and denervation groups (P=NS).

**Table 1.**
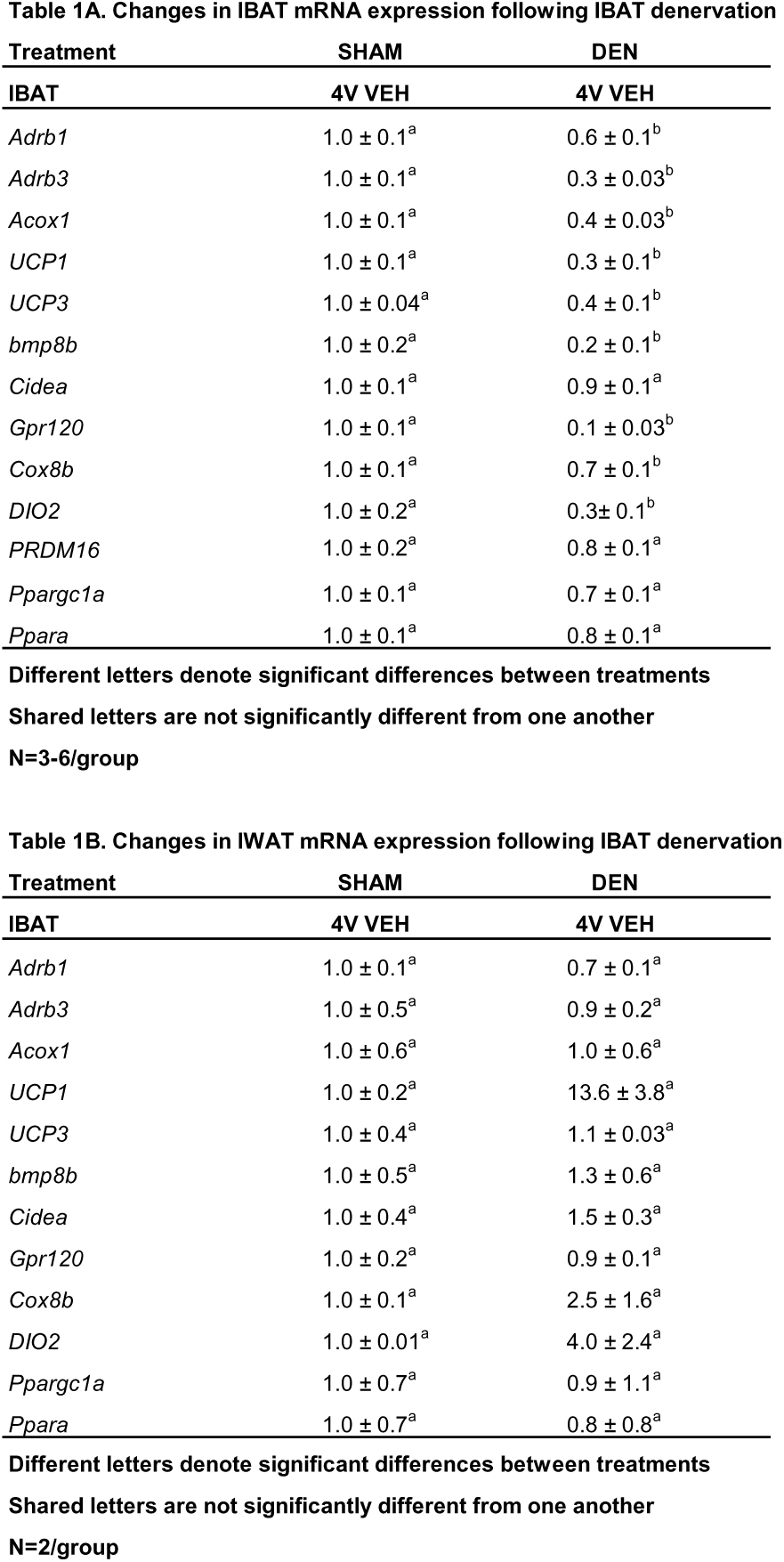
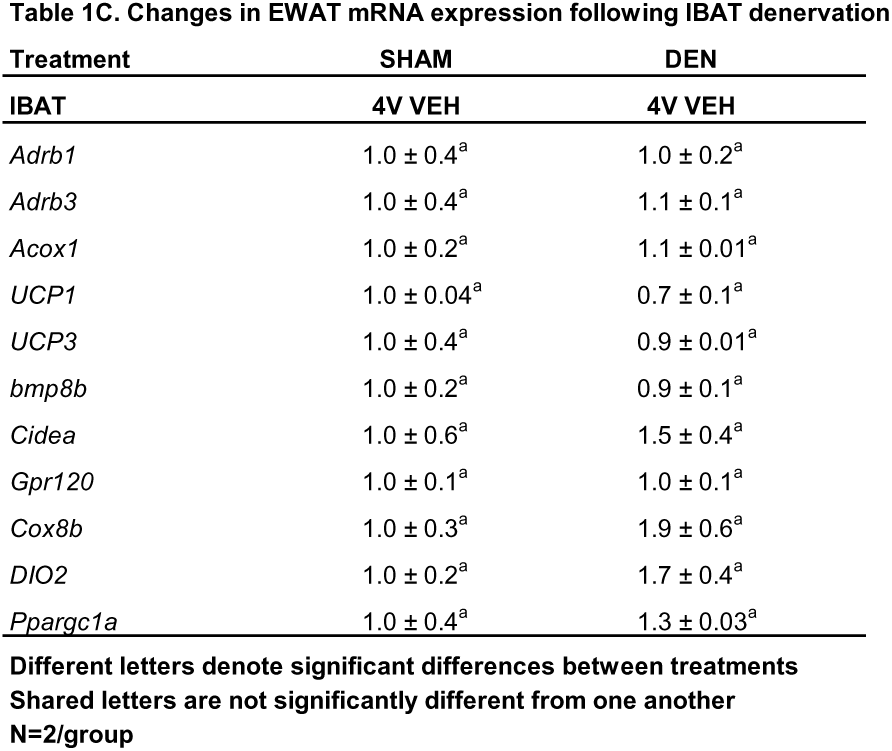
Changes in IBAT, IWAT and EWAT gene expression following sham or IBAT denervation in male DIO mice. *A*, Changes in IBAT mRNA expression following IBAT denervation; *B*, Changes in IWAT mRNA expression following IBAT denervation; *C*, Changes in EWAT mRNA expression following IBAT denervation. Shared letters are not significantly different from one another. Data are expressed as mean ± SEM. \**P*<0.05 OT vs. vehicle (N=2-6/group).

The findings pertaining to IBAT UCP-1 gene expression in surgically denervated mice is consistent with what others have reported with IBAT UCP-1 protein expression from hamsters [51] and mice [52] following chemical (6-OHDA)-induced denervation of IBAT relative to control animals. Similarly, there was a reduction of IBAT UCP-1 mRNA following unilateral or bilateral surgical denervation in hamsters [53] and mice [54; 55], respectively. Similar to our findings, others also found a reduction of IBAT Dio2 and Adrb3 following bilateral surgical denervation in mice [54].

#### IWAT

IWAT UCP-1 [(F(1,2) = 11.131, *P*=0.079)] tended to be elevated in denervated mice relative to sham operated mice (**Table 1B**). There were no significant differences in the mRNAs of any of the other thermogenic markers (*P*=NS; **Table 1B**).

#### EWAT

There tended to be a reduction of UCP-1 in denervated mice compared to sham operated mice [(F(1,2) = 7.000, *P*=0.118)] (**Table 1C**). There were no significant differences in the mRNA levels of any of the other thermogenic genes (*P*=NS; **Table 1C**).

### Study 2: Determine the success of the surgical denervation procedure at 1, 6 and 7-weeks post-sham or denervation in DIO mice

The goal of this study was to 1) verify the success of the SNS denervation procedure in DIO mice by confirming a reduction of NE content that was specific to IBAT relative to other tissues (IWAT, liver and pancreas) and 2) confirm that these changes would persist for 7 weeks. By design, DIO mice were obese as determined by both body weight (45.1±1.1 g) and adiposity (14.9±0.8 g fat mass; 32.9±1.3% adiposity) after maintenance on the HFD for approximately 4.5 months prior to sham/denervation procedures. Sham and denervation groups were matched for body weight, fat mass and lean mass such that there were no differences in baseline body weight, lean mass or fat mass between groups prior to surgery (*P*=NS).

IBAT NE content was reduced by 95.9±2.0, 77.4±12.7 and 93.6±4.6% at 1-[(F(1,8) = 19.636, *P*=0.002)], 6-[(F(1,8) = 20.532, *P*=0.002)] and 7-weeks [(F(1,8) = 34.586, *P*<0.0001)] post-denervation (**Figure 2**) relative to IBAT from sham operated mice. In contrast, NE content was unchanged in IWAT, liver or pancreas in denervated mice relative to sham mice (*P*=NS).

**Figure 2:**
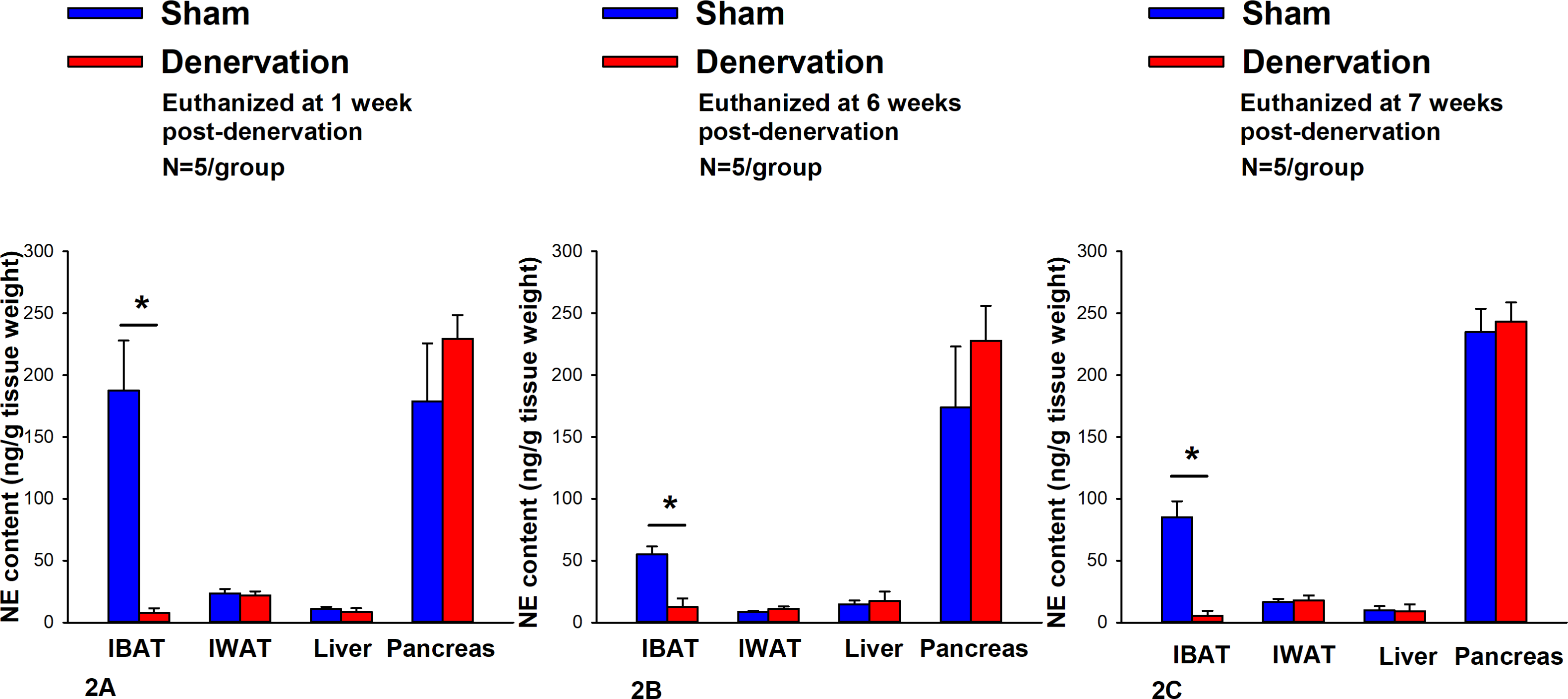
Effect of IBAT surgical denervation procedure on IBAT NE content at 1-, 6- and 7-weeks post-sham or IBAT denervation in DIO mice. Mice were maintained on a HFD (60% kcal from fat) for approximately 4.25 months prior to undergoing a sham (N=5/group) or bilateral surgical IBAT denervation (N=5/group) procedures and were euthanized at 1-, 6- and 7-weeks post-sham (N=15/group) or denervation procedure (N=15/group). NE content was measured from IBAT, IWAT, liver and pancreas. Data are expressed as mean ± SEM. \**P*<0.05, †0.05<*P*<0.1 denervation vs sham.

There were no significant differences in baseline body weights [(F(5,24) = 0.322, *P*=NS), fat mass [(F(5,24) = 0.106, *P*=NS), or lean mass [(F(5,24) = 0.304, *P*=NS) between sham and denervation groups. In addition, there were also no significant differences in body weights between sham and denervation groups at 1-[(F(1,8) = 0.353, *P*=NS), 6-[(F(1,8) = 1.230, *P*=NS) and 7-weeks [(F(1,8) = 0.109, *P*=NS) post-surgeries. In addition, there were also no significant differences in fat mass at 1-[(F(1,8) = 0.353, *P*=NS), 6-[(F(1,8) = 1.230, *P*=NS) and 7-weeks [(F(1,8) = 0.109, *P*=NS) post-denervation relative to sham operated mice. Similarly, no differences were found in lean mass at 1-[(F(1,8) = 0.141, *P*=NS)], 6-[(F(1,8) = 0.015, *P*=NS)] and 7-weeks [(F(1,8) = 0.009, *P*=NS)] post-denervation relative to sham operated mice. These findings are in agreement with others who have reported no difference in body weight [51; 52; 54; 55; 56] or fat mass [52; 55] in hamsters or mice following bilateral surgical or chemical denervation of IBAT.

### Study 3: Determine if SNS innervation of IBAT is reduced in age-matched obese mice relative to lean mice

The goal of this study was to determine if SNS innervation to IBAT is reduced in the obese state relative to lean animals. Given that SNS activation of BAT and BAT activity decline with age ([57]; for review see [58]), DIO mice were age-matched to lean chow-fed control mice. By design, DIO mice were obese as determined by both body weight (46.4±1.2 g) and adiposity (17.8±0.7 g fat mass; 38.3±0.8% adiposity) relative to age-matched chow-fed control mice (33.1±0.4 g) and adiposity (6.3±0.4 g fat mass; 18.9±1.1% adiposity) after maintenance on the HFD and chow, respectively, for approximately 4.5 months prior to sham/denervation procedures.

IBAT NE content was reduced by 38.9±9.8% relative in lean mice relative to age-matched DIO mice [(F(1,8) = 10.757, *P*=0.011)] (**Figure 3**). In addition, NE content in IWAT [(F(1,8) = 19.363, *P*=0.002)] and liver [(F(1,8) = 14.516, *P*=0.005)] was reduced in lean mice compared to age-matched DIO mice. NE content in pancreas also appeared to be reduced in lean mice relative to DIO mice [(F(1,8) = 4.233, *P*=0.074)] while there was no significant difference in NE from EWAT (*P*=NS).

**Figure 3:**
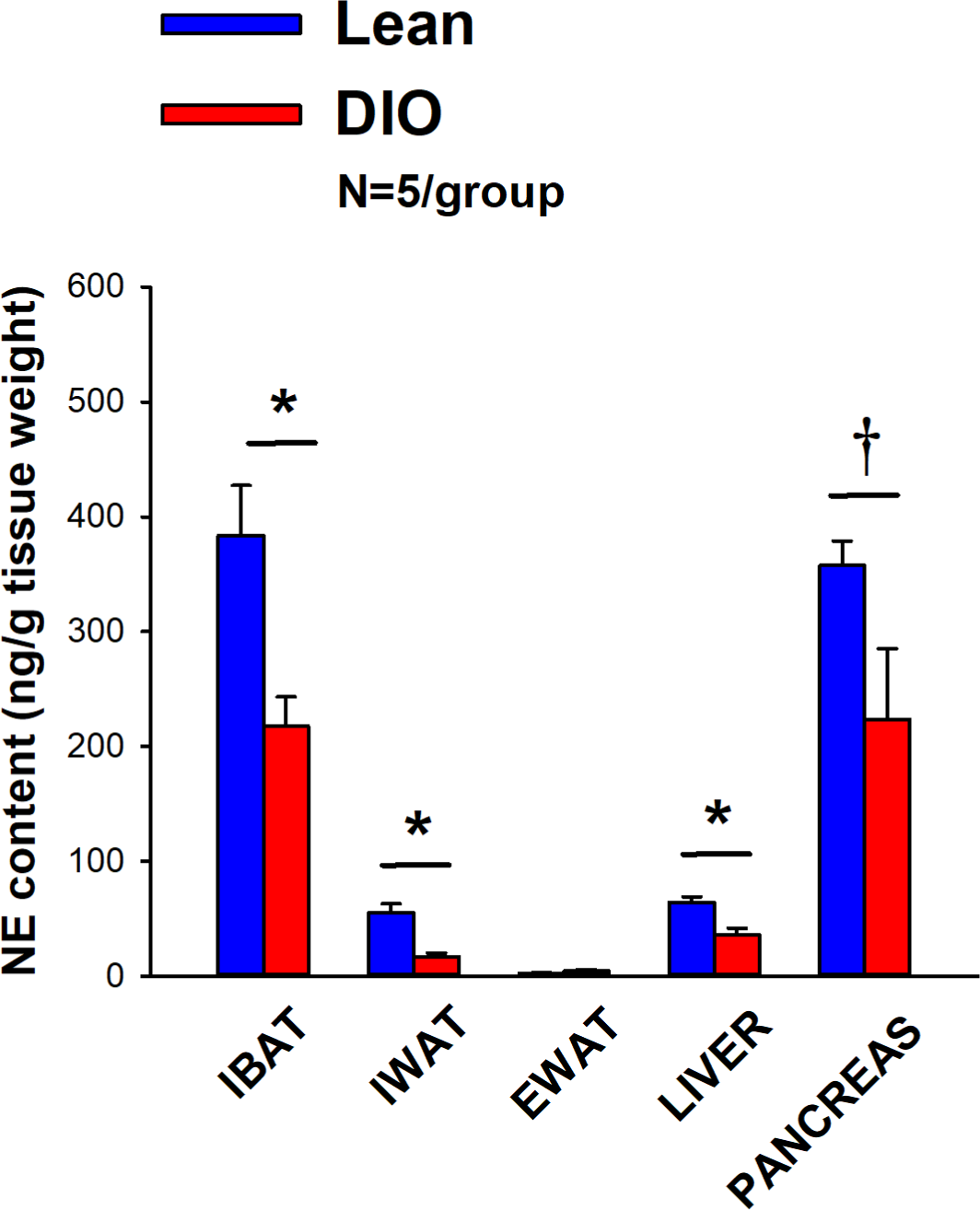
Effect of diet-induced obesity (DIO) on SNS Innervation (NE content) of IBAT, IWAT, EWAT, liver and pancreas at 1-week post-sham in age-matched lean and DIO mice. Age-matched mice were maintained on either chow (16% kcal from fat) or HFD (60% kcal from fat) for approximately 4.5 months prior to undergoing a sham denervation (N=5/group) procedure and were euthanized at 1-week post-sham (N=5/group). NE content was measured from IBAT, IWAT, EWAT, liver and pancreas. Data are expressed as mean ± SEM. \**P*<0.05, †0.05<*P*<0.1 lean vs DIO.

### Study 4: Determine if surgical denervation of IBAT changes the ability of the beta-3 receptor agonist, CL 316243, to increase T_IBAT_ in DIO mice

The goal of this study was to verity that there was no change in the ability of IBAT to respond to direct beta-3 receptor stimulation as a result of the denervation procedure relative to sham operated animals. By design, DIO mice were obese as determined by both body weight (44.2±0.6 g) and adiposity (13.9±0.8 g fat mass; 31.2±1.4% adiposity) after maintenance on the HFD for approximately 4.5 months prior to sham/denervation procedures.

In sham mice, CL 316243 (1 mg/kg) increased T_IBAT_ at 0.25, 0.75, 1, 2, 3 and 4-h post-injection. The lowest dose (0.1 mg/kg) also stimulated T_IBAT_ at 0.25, 0.75, 1, 2, 3 and 4-h post-injection (*P*<0.05; **Figure 4A**). In addition, similar findings were apparent when measuring change in T_IBAT_ relative to baseline T_IBAT_ (**Figure 4B**). CL 316243 also stimulated T_IBAT_ and T_IBAT_ relative to baseline at 24-h post-injection at the high dose (*P*<0.05; data not shown).

**Figure 4.**
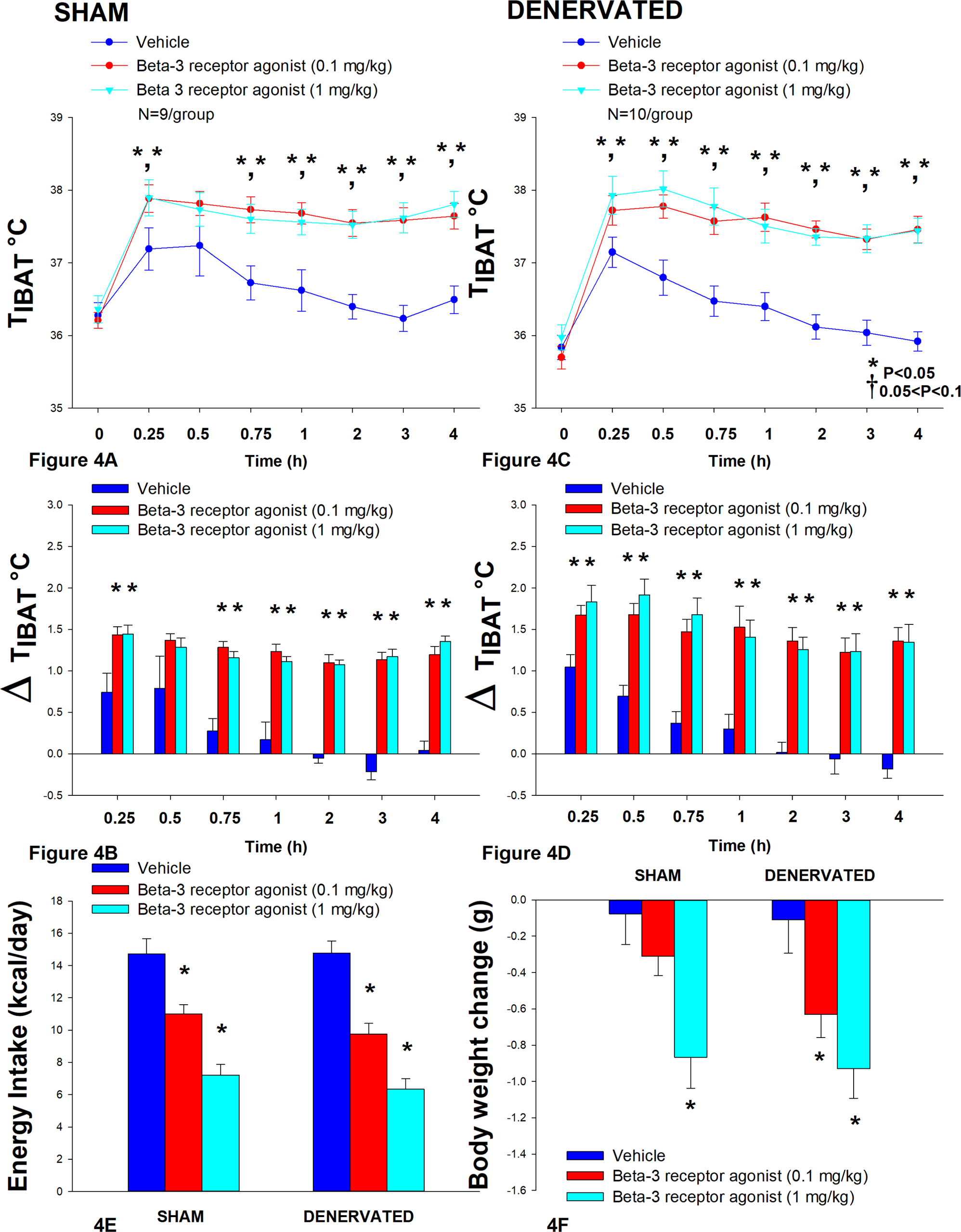
A-F: Effect of systemic beta-3 receptor agonist (CL 31643) administration (0.1 and 1 mg/kg) on IBAT temperature (T_IBAT_), energy intake and body weight post-sham or IBAT denervation in DIO mice. Mice were maintained on HFD (60% kcal from fat; N=9-10/group) for approximately 4.5 months prior to undergoing a sham or bilateral surgical IBAT denervation and implantation of temperature transponders underneath IBAT. Animals were subsequently adapted to a 4-h fast prior to receiving IP injections of CL 316243 (0.1 or 1 mg/kg, IP) or vehicle (sterile water) where each animal received each treatment at approximately 7-day intervals. *A/C*, Effect of CL 316243 on T_IBAT_ in *A)* sham operated or *C)* IBAT denervated DIO mice; *B/D*, Effect of CL 316243 on change in T_IBAT_ relative to baseline T_IBAT_ (delta T_IBAT_) in *B)* sham operated or *D)* IBAT denervated DIO mice; *E*, Effect of CL 316243 on change in energy intake in sham or IBAT denervated DIO mice; *F*, Effect of CL 316243 on change in body weight in sham or IBAT denervated DIO mice. Data are expressed as mean ± SEM. \**P*<0.05, †0.05<*P*<0.1 CL 316243 vs. vehicle.

IBAT NE content was reduced in denervated mice by 97.8±0.5% in denervated mice relative to sham-operated control mice [(F(1,17) = 73.270, *P*<0.001). In contrast, there was no difference in NE content in IWAT, EWAT, liver or pancreas between sham or denervation groups (*P*=NS).

In denervated mice, CL 316243 (1 mg/kg) increased T_IBAT_ at 0.25, 0.5, 0.75, 1, 1.25, 1.5, 1.75, 2, 3 and 4-h post-injection. The lowest dose (0.1 mg/kg) also stimulated T_IBAT_ at 0.25, 0.5, 0.75, 1, 2, 3 and 4-h post-injection ((*P*<0.05; **Figure 4C**). CL 316243 also stimulated T_IBAT_ at 24-h post-injection at the high dose (*P*<0.05; data not shown). Similar findings were apparent when measuring change in T_IBAT_ relative to baseline T_IBAT_ through 240-min post-injection (**Figure 4D**). CL 316243 (1 mg/kg) also tended to stimulate the change in T_IBAT_ relative to baseline T_IBAT_ at 24-h post-injection (*P*=0.050).

Importantly, there was no difference in the T_IBAT_ response to CL 316243 (0.1 or 1 mg/kg) when the data were averaged over the 1-h or 4-h post-injection period between sham and denervated mice (*P*=NS).

Overall, these findings indicate that IBAT denervation did not change the ability of CL 316243 to increase BAT thermogenesis (surrogate measure of EE) in DIO mice relative to sham operated mice.

### Energy intake

In sham mice, CL 316243 reduced daily energy intake at both 0.1 and 1 mg/kg by 25.3 and 51% (P<0.05). Similarly, in denervated rats, CL 316243 also reduced daily energy intake at both 0.1 and 1 mg/kg (*P*<0.05) by 34 and 57.1% relative to vehicle (**Figure 4E**).

### Body weight

CL 316243 did not reduce overall body weight in either group. However, the high dose reduced body weight gain in both sham and denervated mice (*P*<0.05; **Figure 4F**) while the low dose also reduced body weight gain in the denervated mice (*P*<0.05; **Figure 4F**).

Importantly, there was no difference in the effectiveness of CL 316243 (0.1 or 1 mg/kg) to reduce food intake or weight gain between sham and denervated mice (*P*=NS).

However, there appeared to be a more enhanced effect of CL 316243 (0.1 mg/kg) to reduce weight gain (*P*=0.091) in the denervated group relative to the sham group.

Overall, these findings indicate that IBAT denervation did not change the ability of CL 316243 to reduce food intake or body weight gain in DIO mice relative to sham operated mice.

Similar results were also found in lean mice (data not shown). The sympathomimetic, icilin, which may act indirectly stimulate BAT thermogenesis through SNS outflow to IBAT, increased T_IBAT_ (*P*<0.05) throughout the measurement period in both sham and IBAT denervated mice (*P*<0.05; data not shown).

The effects of the sympathomimetic, tyramine (**Supplemental Study 1**), was also examined on T_IBAT_ in DIO mice with intact or impaired SNS innervation of IBAT. Tyramine (19.2 mg/kg) increased T_IBAT_ during the initial part of the measurement period in both sham and IBAT denervated mice (*P*<0.05; **Supplemental Figure 1A-D**).

There was no difference in the T_IBAT_ response to tyramine (19.2 mg/kg) when the data were averaged over the 0.25-h post-injection period between sham and denervated mice (*P*=NS). Similar findings were also observed when comparing T_IBAT_ or change from baseline T_IBAT_ at 0.25-h and 0.5-h post-injection between sham and denervated mice (*P*=NS).

Overall, these findings demonstrate there is not a functional change in the ability of IBAT to respond to direct beta-3 receptor stimulation or other agents, including icilin, that may activate BAT through a direct non-beta-3 receptor mechanism in denervated mice relative to sham animals.

### Study 5: Determine the extent to which OT-induced activation of sympathetic outflow to IBAT contributes to its ability to increase T_IBAT_ in DIO mice

After having confirmed there was no functional defect in the ability of IBAT to respond to direct beta-3 receptor stimulation (**Study 3**), the goal of this study was to determine if OT-elicited elevation of T_IBAT_ requires intact SNS outflow to IBAT. By design, DIO mice were obese as determined by both body weight (43.6±0.7 g) and adiposity (13.8±0.6 g fat mass; 31.5±1.1% adiposity) after maintenance on the HFD for approximately 4.25 months prior to sham/denervation procedures. Note that the T_IBAT_ data from sham-operated DIO mice has been previously published [22].

IBAT NE content was reduced by 94.9±3.3% in denervated mice relative to sham-operated control mice [(F(1,19) = 33.427, *P*<0.001). In contrast, there was no difference in NE content in IWAT, EWAT, liver or pancreas between sham or denervation groups (*P*=NS).

In sham mice, 4V OT (5 μg) increased T_IBAT_ at 1 and 1.25-h post-injection (P<0.05) and tended to stimulate T_IBAT_ at 0.75, 1.5 and 1.75-h post-injection (0.05<*P*<0.1). The lowest dose (1 μg) also stimulated T_IBAT_ at 0.75 and 1-h post-injection and tended to stimulate T_IBAT_ at 1.25-h post-injection (0.05<*P*<0.1; **Figure 5A**). Similar findings were apparent when measuring change in T_IBAT_ relative to baseline T_IBAT_ (**Figure 5B**).

**Figure 5.**
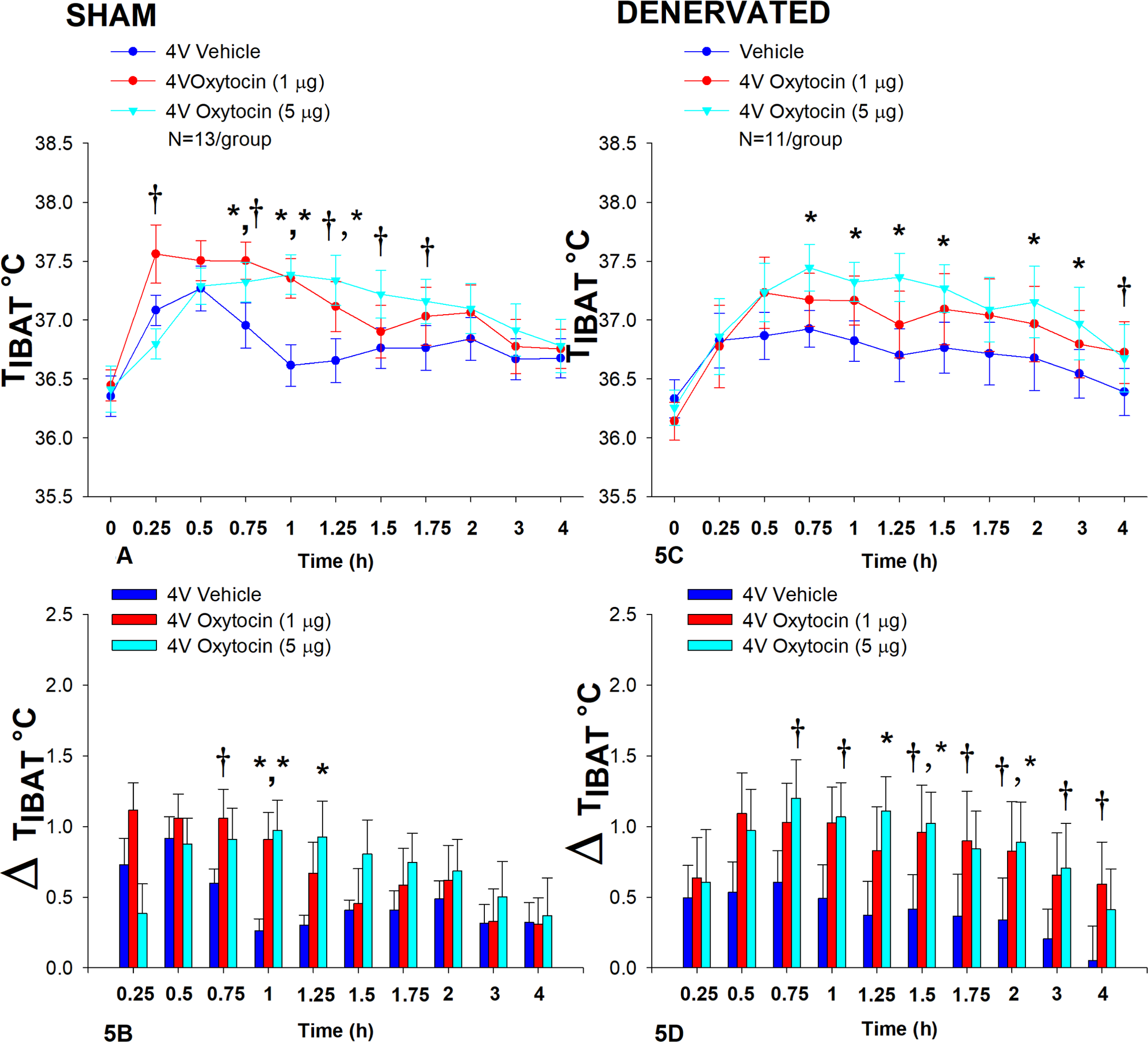
A-D: Effect of acute 4V OT administration (1 and 5 μg) on T_IBAT_ post-sham or IBAT denervation in male DIO mice. Mice were maintained on HFD (60% kcal from fat; N=11-13/group) for approximately 4.25 months prior to undergoing a sham or bilateral surgical IBAT denervation and implantation of temperature transponders underneath IBAT. Mice were subsequently implanted with 4V cannulas and allowed to recover for 2 weeks prior to receiving acute 4V injections of OT or vehicle. Animals were subsequently adapted to a 4-h fast prior to receiving acute 4V injections of OT or vehicle *A/C*, Effect of acute 4V OT on T_IBAT_ in *A)* sham operated or *C)* IBAT denervated DIO mice; *B/D*, Effect of acute 4V OT on change in T_IBAT_ relative to baseline T_IBA_T (delta T_IBAT_) in *B)* sham operated or *D)* IBAT denervated DIO mice; Data are expressed as mean ± SEM. \**P*<0.05, †0.05<*P*<0.1 OT vs. vehicle.

In denervated mice, OT (5 μg) increased T_IBAT_ at 0.75, 1, 1.25, 1.5, 2 and 3-h post-injection (P<0.05). The lowest dose (1 μg) tended to stimulate T_IBAT_ at 4-h and 24-h post-injection (0.05<*P*<0.1; **Figure 5C**). Similar findings were apparent when measuring change in T_IBAT_ relative to baseline T_IBAT_ (**Figure 5D**). In addition, 4V OT elevated T_IBAT_ in lean denervated mice (data not shown).

Overall, these findings demonstrate that SNS innervation of IBAT is not a predominant mediator of OT-elicited elevations of BAT thermogenesis.

In contrast to the pattern observed following 4V administration, systemic administration of OT produced an initial reduction of T_IBAT_ (*P*<0.05) followed by an increase in T_IBAT_ (*P*<0.05; **Supplemental Figure 3A-D**). Similar to OT, the melanocortin 3/4 receptor agonist, melanotan II (MTII), produced an initial reduction of T_IBAT_ (*P*<0.05; data not shown) followed by a subsequent increase in T_IBAT_ (data not shown; *P*<0.05).

### Study 6A: Determine the extent to which sympathetic outflow to IBAT contributes to the ability of OT to elicit weight loss in DIO mice (0927.27)

The goal of this study was to determine if OT-elicited weight loss requires intact SNS outflow to IBAT. By design, DIO mice were obese as determined by both body weight (47.9±0.6 g) and adiposity (17.7±0.5 g fat mass; 36.9±0.7% adiposity) after maintenance on the HFD for approximately 4.25-4.5 months prior to sham/denervation procedures.

IBAT NE content was reduced in denervated mice by 93.9±2.3% in denervated mice relative to sham-operated control mice [(F(1,28) = 23.306, *P*=0.000). In contrast, there was no difference in NE content in IWAT, EWAT, liver or pancreas between sham or denervation groups (*P*=NS)

Chronic 4V OT reduced body weight by 5.7±2.23% and 6.6±1.4% in sham and denervated mice (*P*<0.05), respectively, and this effect was similar between groups (*P*=NS). OT produced corresponding reductions in fat mass (*P*<0.05) and this effect was also similar between groups (*P*=NS).

As expected, 4V vehicle resulted in 7.5±3.0% weight gain relative to vehicle pre-treatment [(F(1,8) = 5.677, *P*=0.044)]. In contrast, 4V OT reduced body weight by 5.7±2.23% relative to OT pre-treatment [(F(1,7) = 5.903, *P*=0.045)] (**Figure 6A**) and reduced weight gain (**Figure 6B**) between days 4-29 of the infusion period (*P*<0.05). 4V OT tended to reduce weight gain between days 2-3 (0.05<*P*<0.1). OT produced a corresponding reduction in fat mass [(F(1,13) = 5.190, *P*=0.040)] (**Figure 6C**) with no effect on lean body mass (*P*=NS). These effects were mediated, at least in part, by a modest reduction of energy intake that tended to be present during weeks 1 (*P*=0.067; **Figure 6D**) and 2 (*P*=0.092).

**Figure 6.**
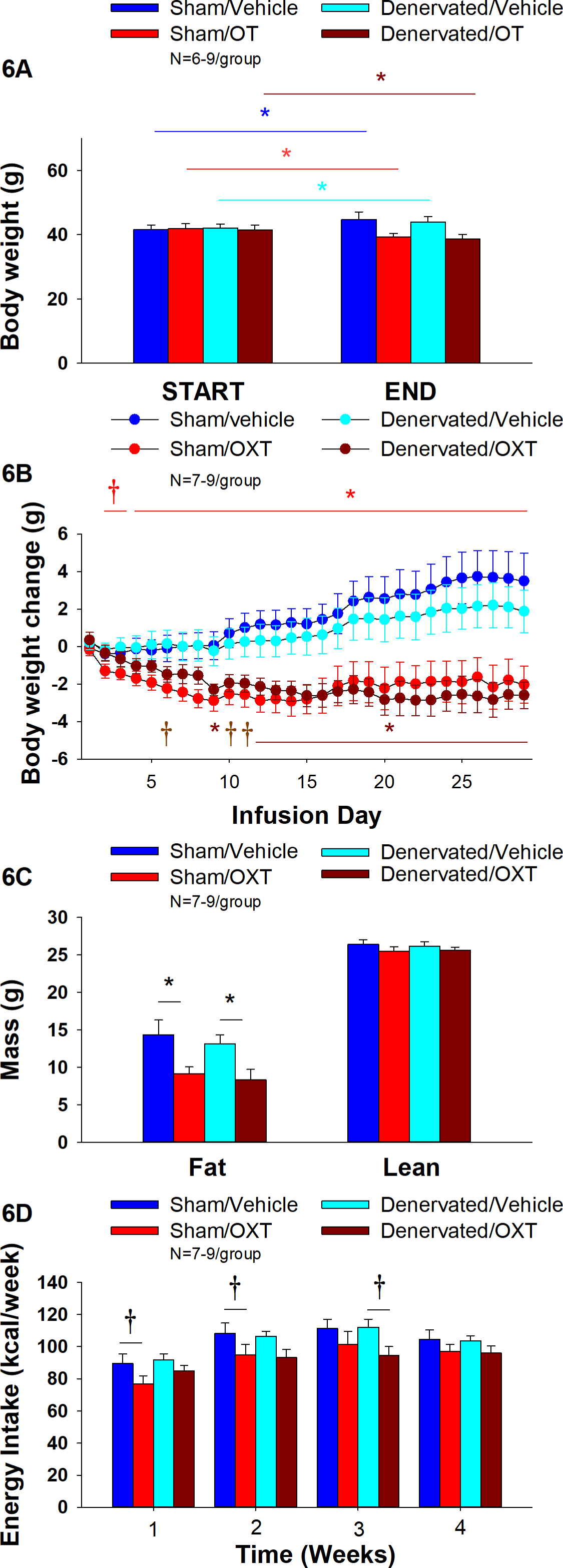
A-D: Effect of chronic 4V OT infusions (16 nmol/day) on body weight, adiposity and energy intake post-sham or IBAT denervation in male DIO mice. *A*, Mice were maintained on HFD (60% kcal from fat; N=7-9/group) for approximately 4.25-4.5 months prior to undergoing a sham or bilateral surgical IBAT denervation. Mice were subsequently implanted with 4V cannulas and allowed to recover for 2 weeks prior to being implanted with subcutaneous minipumps that were subsequently attached to the 4V cannula. *A*, Effect of chronic 4V OT or vehicle on body weight in sham operated or IBAT denervated DIO mice; *B*, Effect of chronic 4V OT or vehicle on body weight change in sham operated or IBAT denervated DIO mice; *C*, Effect of chronic 4V OT or vehicle on adiposity in sham operated or IBAT denervated DIO mice; *D*, Effect of chronic 4V OT or vehicle on adiposity in sham operated or IBAT denervated DIO mice. Data are expressed as mean ± SEM. \**P*<0.05, †0.05<*P*<0.1 OT vs. vehicle.

In denervated mice, 4V vehicle treatment resulted in 4.3±2.6% weight gain relative to vehicle pre-treatment [(F(1,5) = 17.371, *P*=0.009)]. In contrast, 4V OT reduced body weight by 6.6±1.4% relative to OT pre-treatment [(F(1,5) = 15.883, *P*=0.010)] (**Figure 6A**) and it reduced weight gain (**Figure 6B**) throughout the 29-day infusion period. OT treatment reduced weight gain on days 9 and 12-29 (*P*<0.05) and it tended to reduce weight gain on days 6 (*P*=0.095), 10 (*P*=0.059) and 11 (*P*=0.055). OT produced a corresponding reduction in fat mass [(F(1,11) = 7.101, *P*=0.022)] (**Figure 6C**) with no effect on lean body mass (*P*=NS). These effects were mediated, at least in part, by a modest reduction of energy intake that tended to be present during weeks 2 (*P*=0.147) and 3 (*P*=0.064; **Figure 6D**).

Based on these collective findings, we conclude that SNS innervation of IBAT is not a predominant contributor of oxytocin-elicited increases in BAT thermogenesis (surrogate measure of energy expenditure), weight loss and reduction of fat mass.

### Adipocyte size

#### Sham

There was a significant effect of 4V OT to reduce EWAT adipocyte size in sham operated mice [(F(1,13) = 5.729, *P*=0.032)] (**Figure 7B**) while it had no significant effect on IWAT adipocyte size (*P*=NS) (**Figure 7A**).

**Figure 7.**
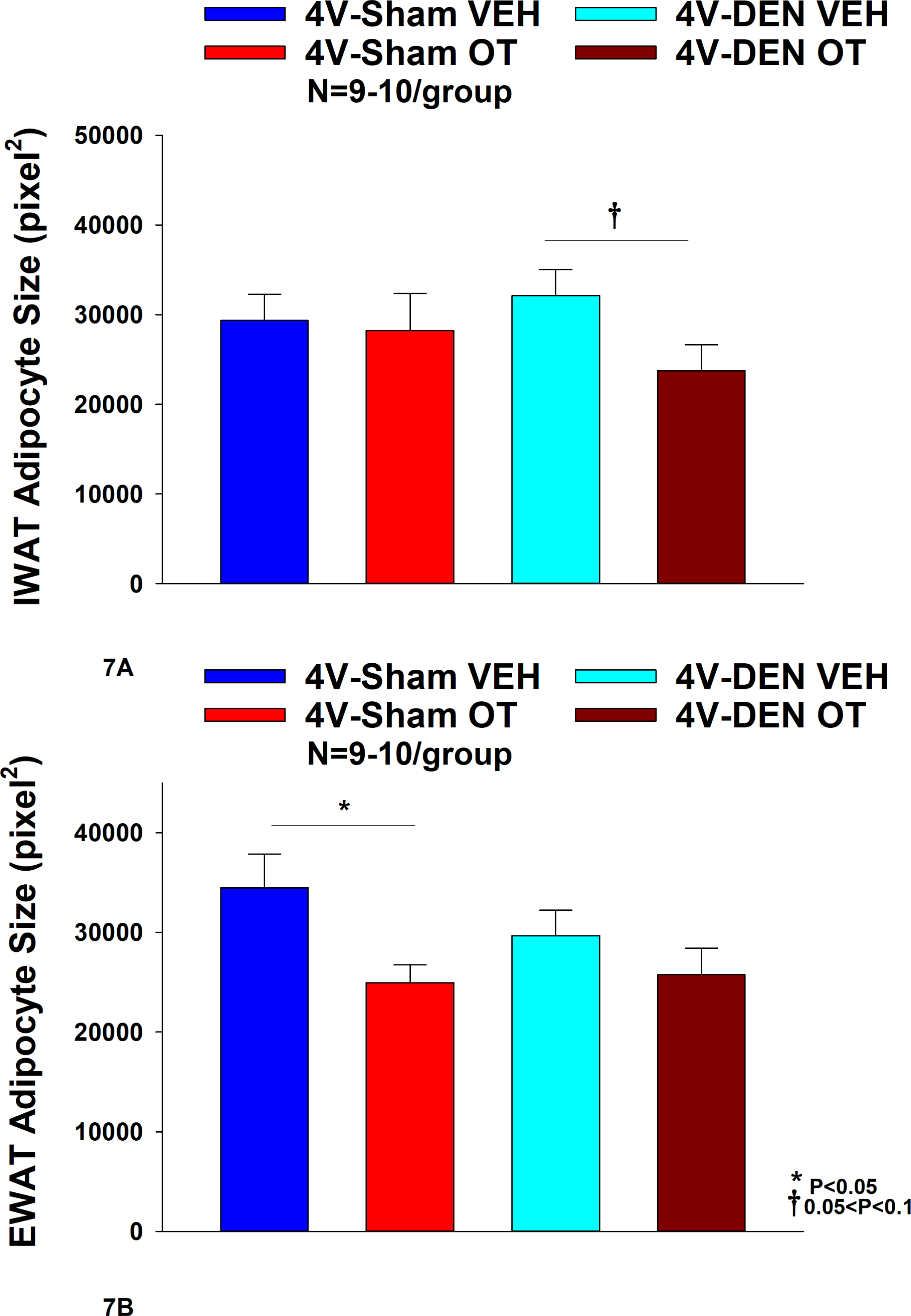
A-B: Effect of chronic 4V OT infusions (16 nmol/day) on adipocyte size post-sham or IBAT denervation in male DIO mice. *A*, Adipocyte size (pixel^2^) was measured in IWAT from mice that received chronic 4V infusion of OT (16 nmol/day) or vehicle in sham or IBAT denervated DIO mice (N=9-10/group). *B*, Adipocyte size was measured in EWAT from rats that received chronic 4V infusion of OT (16 nmol/day) or vehicle in sham operated or IBAT denervated mice (N=9-10/group). Data are expressed as mean ± SEM. \**P*<0.05 OT vs. vehicle.

#### Denervation

In contrast to the effects observed in sham mice, 4V OT had no significant effect on EWAT adipocyte size in IBAT denervated mice (*P*=NS) (**Figure 7B**). However, there was a tendency of 4V OT to reduce IWAT adipocyte size in IBAT denervated mice [(F(1,11) = 4.233, *P*=0.064)] (**Figure 7A**).

### Plasma Hormone Concentrations

To characterize the endocrine and metabolic effects between sham and denervated DIO mice, we measured blood glucose levels and plasma concentrations of leptin, insulin, FGF-21, irisin, adiponectin, FFA, free glycerol (FG) and total cholesterol (TC) post-sham and denervation procedure. We found an overall effect of 4V OT to reduce plasma leptin [(F(3,26) = 3.839, *P*=0.021)]. Specifically, 4V OT treatment was associated with a reduction of plasma leptin in the sham group (*P*<0.05; **Table 2**) which coincided with OT-elicited reductions in fat mass. We also found that 4V OT treatment tended to reduce plasma insulin in the denervation group relative to the 4V vehicle sham group (*P*=0.081; **Table 2**). In addition, there was an overall effect of OT to reduce plasma total cholesterol [(F(3,26) = 5.806, *P*=0.004)]. 4V OT treatment was associated with a significant reduction of total cholesterol in both groups (*P*<0.05; **Table 2**).

**Table 2.**
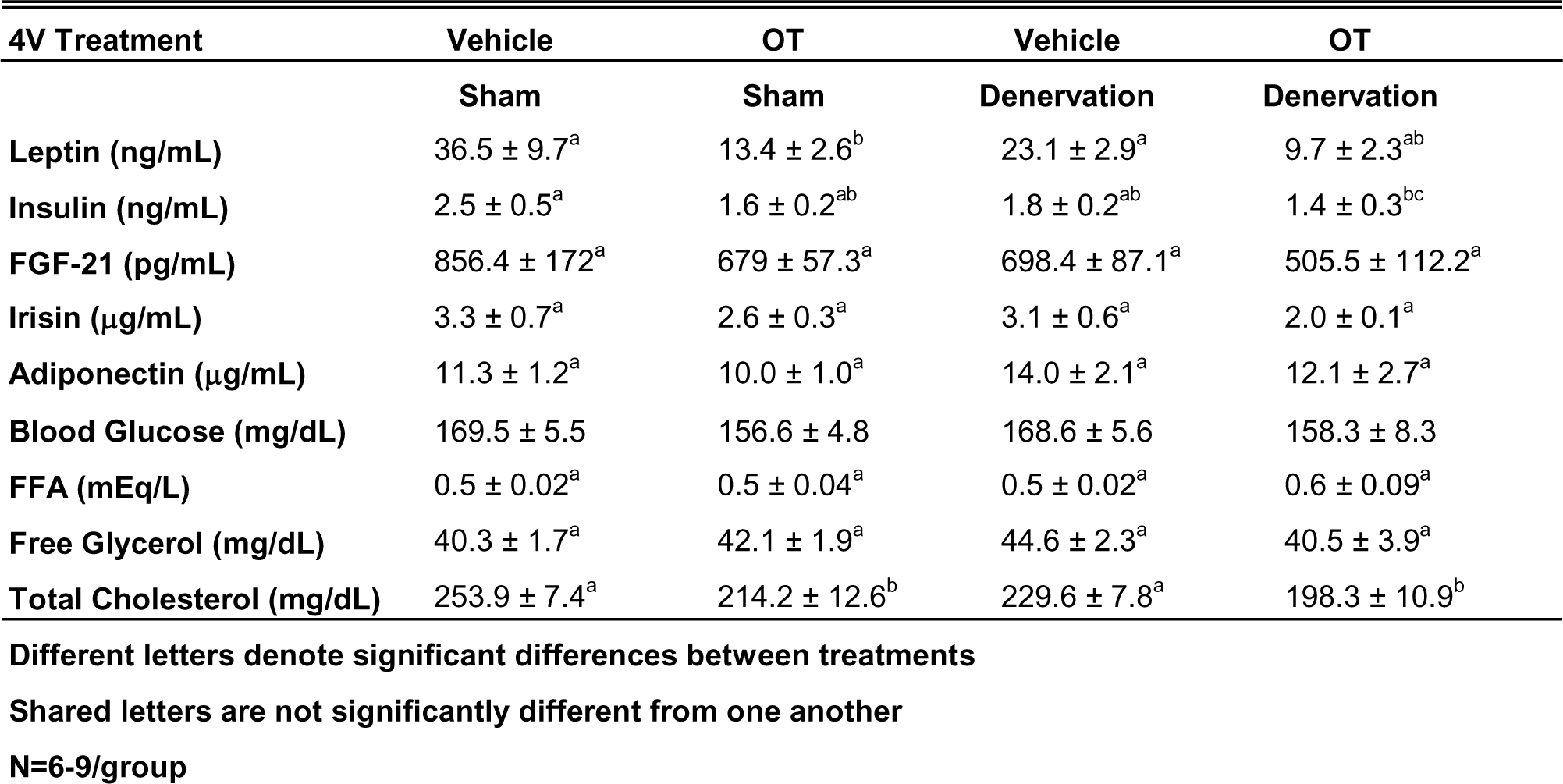
Plasma measurements following 4V infusions of OT (16 nmol/day) or vehicle in sham and IBAT denervated diet-induced obese mice. Data are expressed as mean ± SEM. \**P*<0.05 OT vs. vehicle (N=6-9/group).

### Study 6B: Determine the extent to which sympathetic outflow to IBAT contributes to the ability of OT to impact thermogenic gene expression in IBAT, IWAT and EWAT in DIO mice

#### IBAT

There was a reduction of IBAT UCP-1 [(F(1,7) = 38.1, *P*<0.001)], DIO2 [(F(1,7) = 12.669, *P*=0.009)], Gpr120 [(F(1,7) = 65.965, *P*<0.001)], Adrb3 [(F(1,7) = 65.916, *P*=0.000)], Adrb1 [(F(1,7) = 8.015, *P*=0.025)], Acox1 [(F(1,7) = 58.261, *P*<0.001)], bmp8b [(F(1,7) = 19.636, *P*=0.003)], cox8b [(F(1,6) = 16.403, *P*=0.007)] and UCP-3 mRNA expression [(F(1,7) = 57.665, *P*<0.001)] in denervated mice relative to IBAT from sham operated mice (*P*<0.05; **Table 3A**). There were no significant differences in IBAT CIDEA, PPARGC1, PPARA and PRDM16 mRNA expression between sham and denervation groups (P=NS).

**Table 3.**
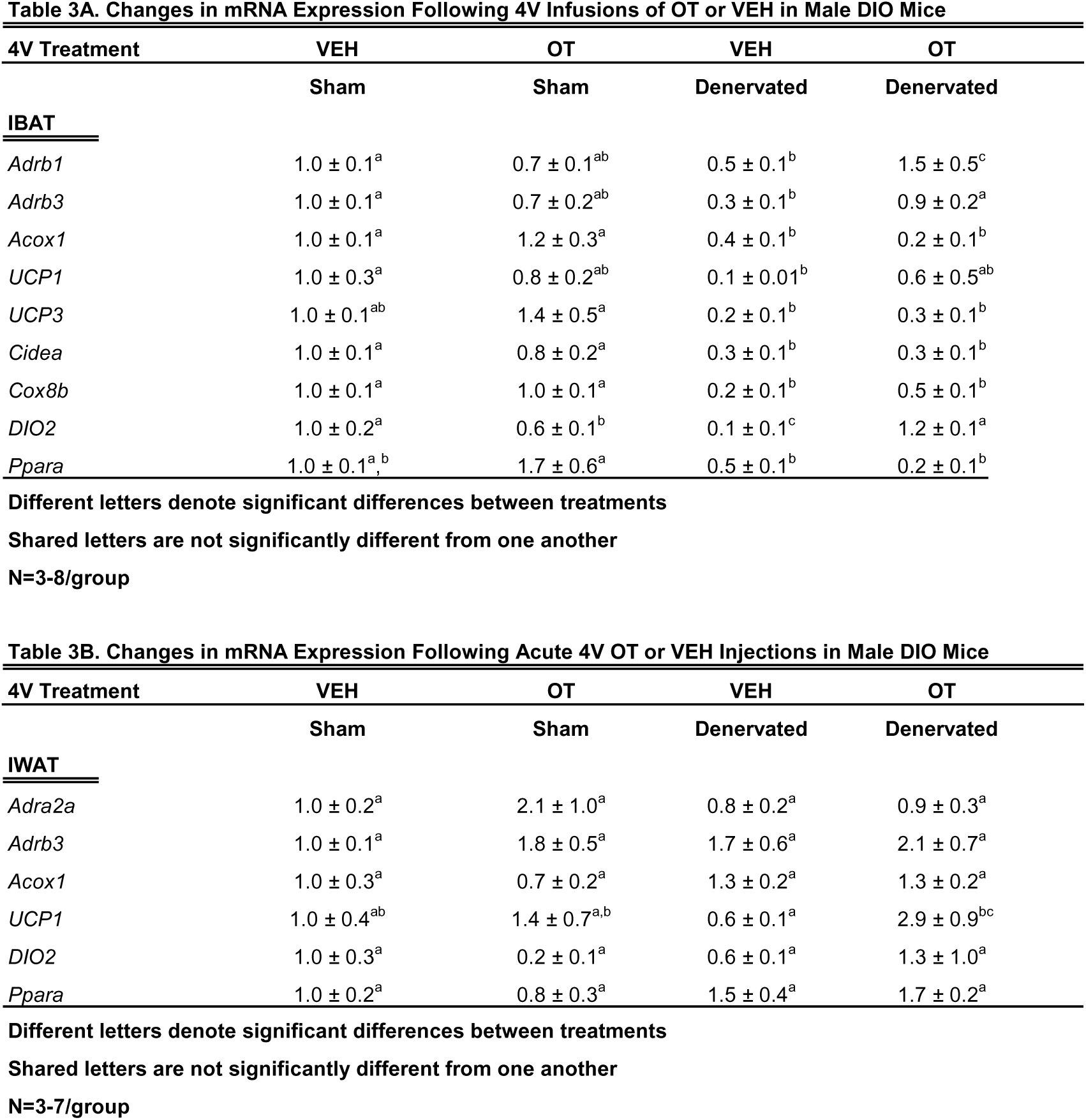
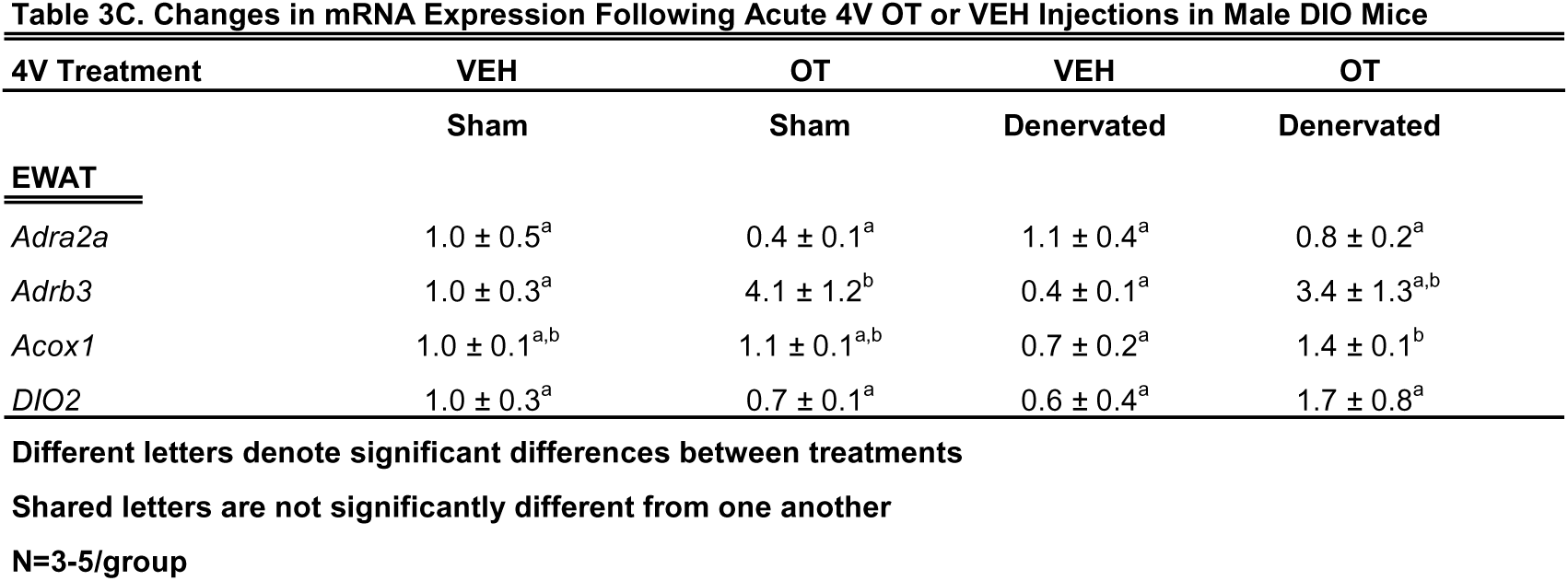
Changes in IBAT, IWAT and EWAT gene expression following 4V infusions of OT or vehicle in male sham or IBAT denervated DIO mice. *A*, Changes in IBAT mRNA expression 4V infusions of OT or vehicle in male sham or IBAT denervated DIO mice; *B*, Changes in IWAT mRNA expression following 4V infusions of OT or vehicle in male sham or IBAT denervated DIO mice; *C*, Changes in EWAT mRNA expression following 4V infusions of OT or vehicle in male sham or IBAT denervated DIO mice. Shared letters are not significantly different from one another. Data are expressed as mean ± SEM. \**P*<0.05 OT vs. vehicle (N=3-8/group).

The findings pertaining to IBAT UCP-1 gene expression in surgically denervated mice is consistent with what others have reported with IBAT UCP-1 protein expression from hamsters [51] and mice [52] following chemical (6-OHDA)-induced denervation of IBAT relative to control animals. Similarly, there was a reduction of IBAT UCP-1 mRNA following unilateral or bilateral surgical denervation in hamsters [53] and mice [54; 55], respectively. Similar to our findings, others also found a reduction of IBAT Dio2 and Adrb3 following bilateral surgical denervation in mice [54]. The findings pertaining to IBAT UCP-1 gene expression in denervated mice is consistent with what others have reported with IBAT UCP-1 protein expression from hamsters [51] and mice [52] with chemical (6-OHDA)-induced denervation of IBAT relative to control animals.

#### IWAT

IWAT UCP-1 [(F(1,2) = 11.131, *P*=0.079)] tended to be elevated in denervated mice relative to sham operated mice (**Table 3B**). There were no significant differences in any of the other thermogenic markers (*P*=NS; **Table 3B**).

#### EWAT

There tended to be a reduction of UCP-1 in denervated mice compared to sham operated mice [(F(1,2) = 7.000, *P*=0.118)] (**Table 3C**). There were no significant differences in any of the other thermogenic markers (*P*=NS; **Table 3C**).

### Study 7: Determine the extent to which systemic (subcutaneous) infusion of a centrally effective dose of OT (16 nmol/day) elicits weight loss in DIO mice

As expected, weight gain of DIO mice increased over the month of vehicle treatment relative to pre-treatment [(F(1,10) = 16.901, *P*=0.002)] (**Figure 8A**). In contrast to the weight lowering effects of 4V OT (16 nmol/day), systemic OT (16 nmol/day) resulted in a significant elevation of body weight relative to OT pre-treatment [(F(1,12) = 11.138, *P*=0.006)] (**Figure 8A**; *P*<0.05). Furthermore, SC OT, at a 3-fold higher dose (50 nmol/day), also resulted in a significant elevation of body weight relative to pre-treatment [(F(1,12) = 10.424, *P*=0.007)]. However, SC OT (16 and 50 nmol/day) was able to reduce weight gain (**Figure 8B**) relative to vehicle treatment throughout the 28-day infusion period. SC OT (50 nmol/day), at a dose that was at least 3-fold higher than the centrally effective dose (16 nmol/day), reduced weight gain throughout the entire 28-day infusion period. SC OT (16 nmol/day) treated mice had reduced weight gain between days 17-28 (*P*<0.05) but did not have the net weight loss seen when this dose was given centrally. SC OT (16 and 50 nmol/day) reduced fat mass [(F(2,35) = 5.558, *P*=0.008)] (**Figure 8C**; *P*<0.05) and produced a corresponding reduction of plasma leptin [(F(2,35) = 3.890, *P*=0.03)], (**Table 4**) with no effect on lean body mass (*P*=NS). These effects that were mediated, at least in part, by a modest reduction of energy intake that was apparent during week 3 of SC OT (50 nmol/day) treatment (**Figure 8D**; *P*<0.05).

**Figure 8.**
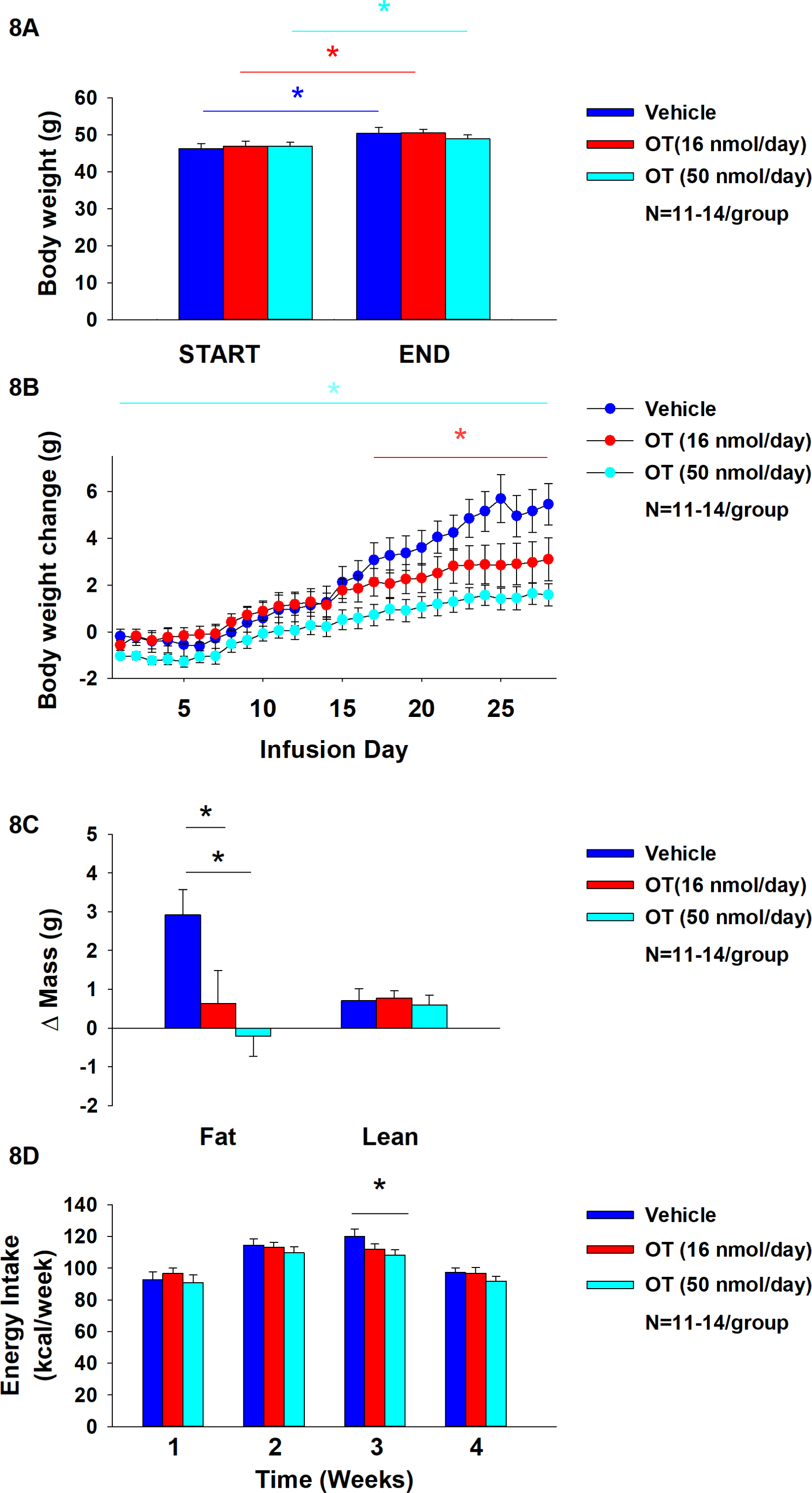
A-D: Effect of chronic subcutaneous OT infusions (16 and 50 nmol/day) on body weight, adiposity and energy intake in male DIO mice. *A*, Mice were maintained on HFD (60% kcal from fat; N=11-14/group) for approximately 4-4.25 months prior to being implanted with temperature transponders and allowed to recover for 1-2 weeks prior to being implanted with subcutaneous minipumps. *A*, Effect of chronic subcutaneous OT or vehicle on body weight in DIO mice; *B*, Effect of chronic subcutaneous OT or vehicle on body weight change in DIO mice; *C*, Effect of chronic subcutaneous OT or vehicle on adiposity in DIO mice; *D*, Effect of chronic subcutaneous OT or vehicle on adiposity in DIO mice. Data are expressed as mean ± SEM. \**P*<0.05, †0.05<*P*<0.1 OT vs. vehicle.

**Table 4.**
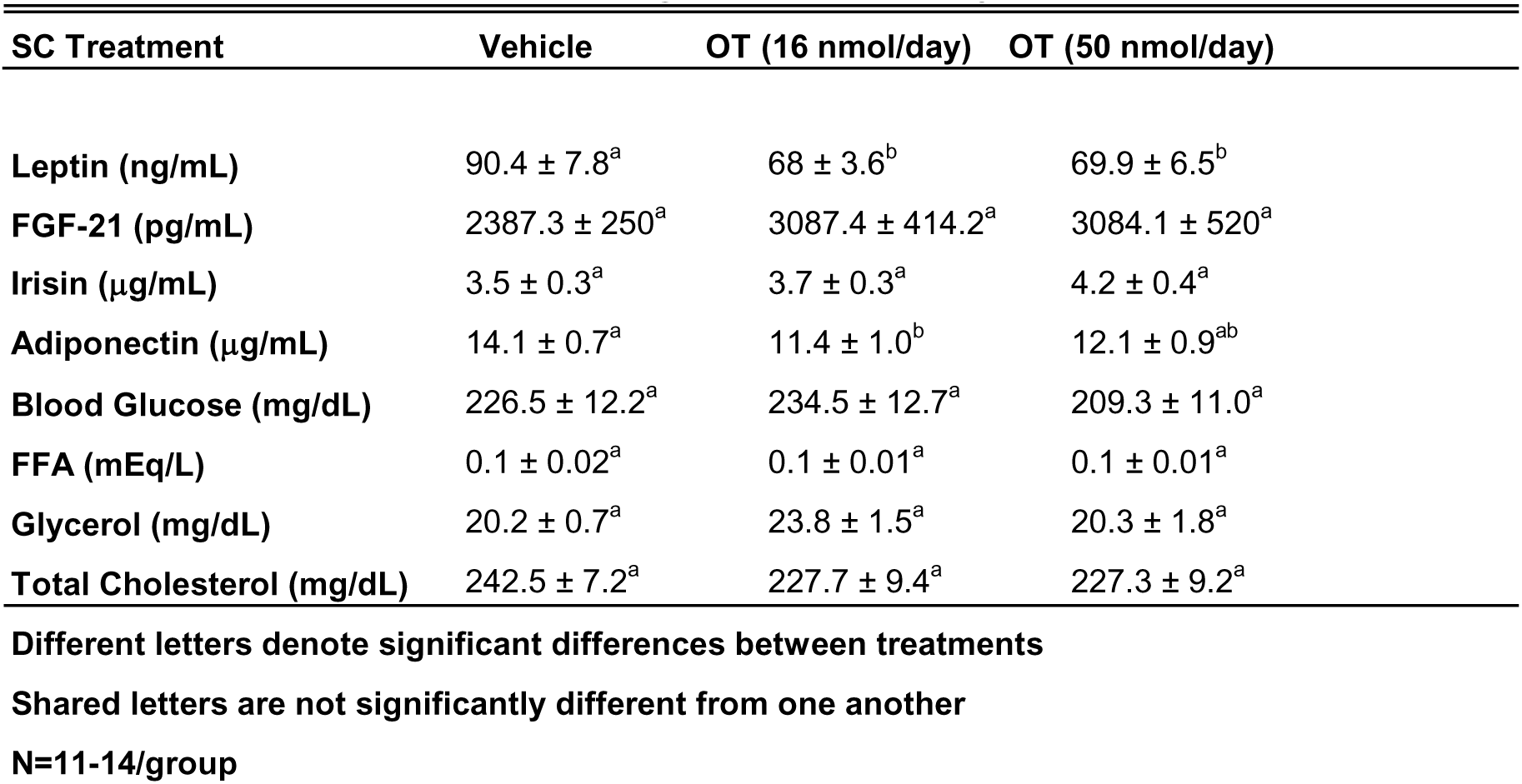
Plasma measurements following systemic infusions of OT (16 and 50 nmol/day) or vehicle in adult male DIO mice. Data.

## Discussion

The goal of the current studies was to determine if sympathetic innervation of IBAT is required for OT to increase non-shivering BAT thermogenesis and reduce body weight and adiposity in male DIO mice. To assess if OT-elicited changes in non-shivering BAT thermogenesis require intact SNS outflow to IBAT, we examined the effects of acute 4V OT (1, 5 μg) on T_IBAT_ in DIO mice following bilateral surgical SNS denervation to IBAT. We found that the high dose (5 µg) elevated T_IBAT_ similarly in sham mice as in denervated mice. We subsequently determined if OT-elicited reductions of body weight and adiposity require intact SNS outflow to IBAT. To accomplish this, we determined the effect of bilateral surgical denervation of IBAT on the ability of chronic 4V OT (16 nmol/day) administration to reduce body weight, adiposity and food intake in DIO mice. We found that chronic 4V OT produced comparable reductions of body weight and adiposity in denervated mice, as well as sham mice (*P*<0.05), supporting the hypothesis that sympathetic innervation of IBAT is not a predominant mediator of OT-elicited increases of non-shivering BAT thermogenesis and reductions of body weight and adiposity in male DIO mice.

Our finding that OT produced these effects when given into the hindbrain (4V) suggest that hindbrain populations and/or spinal cord populations may contribute to the effects of OT-elicited thermogenesis and browning of WAT in mice. Recent findings highlight the presence of both overlapping and nonoverlapping CNS circuits that control SNS outflow to IWAT and IBAT [51]. Namely, parvocellular PVN OT neurons have multi-synaptic projections to IBAT [59; 60], IWAT [59; 61] and EWAT [61; 62]. A small subset of parvocellular PVN OT neurons overlap and project to both IBAT and IWAT [59]. OT neurons are anatomically situated to control SNS outflow to IBAT and IWAT and increase BAT thermogenesis and browning of IWAT, respectively. It is not clear if these effects are mediated by the same OT neurons or through distinct OT neurons that project to the hindbrain nucleus of the solitary tract or nucleus tractus solitarius (NTS) [63; 64] and/or spinal cord [64], both of which are sites that can control SNS outflow to IBAT and BAT thermogenesis [65; 66]. Ong and colleagues recently found that viral-elicited knockdown of OTR mRNA within the dorsal vagal complex was unable to block the effects of 4V OT to increase core temperature [67] suggesting that other OTRs within other areas of the hindbrain and/or spinal cord may contribute to these effects in a rat model. It will be important to determine whether 4V OT-elicited BAT thermogenesis is mediated by OTRs within regions of the NTS not targeted by Ong and colleagues [67], other hindbrain areas such as the raphe pallidus [29; 66; 68; 69; 70] or spinal cord [23].

One outstanding question is how 4V OT is activating IBAT if not through SNS outflow to IBAT. Our findings that 1) systemic administration of OT, at a dose that stimulated T_IBAT_ when given into the 4V, did not fully recapitulate the temporal profile we found following 4V administration and 2) 4V administration failed to reproduce the reduction of T_IBAT_ observed following systemic administration suggest that 4V OT is not likely leaking into the periphery to act at peripheral OTRs. One possibility might be that 4V OT activates OTRs within the hindbrain and/or spinal cord that results in the stimulation of epinephrine from the adrenal gland and subsequent activation of IBAT. Epinephrine has been previously found to stimulate both lipolysis and respiration from brown adipocytes derived from rat IBAT [71] and it has only a 2.5-fold lower affinity for recombinant beta-3 receptors in CHO cells than NE [72]. In addition, mice that lack epinephrine are able to maintain body temperature in response to cold stress but fail to show an increase in IBAT UCP-1 or PGC1-alpha [73] suggesting that non-UCP mechanisms may be involved to increase or retain heat. However, while it is clear that the hindbrain and spinal cord are part of a multi-synaptic projection to the adrenal gland [74; 75], only 1% of PVN OT neurons within either the parvocellular PVN or magnocellular PVN were found to have multi-synaptic projections to the adrenal gland [74]. Whether 4V OT activates a hindbrain or spinal cord projection to the adrenal gland resulting in the release of epinephrine and subsequent activation of IBAT will need to be examined in future studies.

Recent findings also implicate a potentially important role of peripheral OT receptors in the control of body weight in rodents. Similar to our studies, Yuan and colleagues found systemic OT (100 nmol/day) reduced weight gain in high fat diet-fed mice and these effects were associated with decreased adipocyte size (IWAT and EWAT). Asker and colleagues extended these findings and reported that a novel BBB-impermeable OT analog, OT-B12, reduced food intake in rats, thus providing additional evidence for the importance of peripheral OTRs in the control of food intake [76]. These findings are in line with earlier studies by Iwasaki and colleagues who found that the effects of peripheral OT to reduce food intake were attenuated in vagotomized mice [77; 78]. Similarly, previous findings from our lab also indicated that peripheral administration of a non-BBB penetrant OTR antagonist, L-371257, resulted in a modest stimulation of food intake and body weight gain in rats [79]. These findings suggest that, in addition to a central mechanism mediated through hindbrain and/or spinal cord OTRs, OT may also act peripherally to reduce adipocyte size through a direct action on OTRs found on adipocytes [9; 80; 81].

The role of peripheral OTR in the control of BAT thermogenesis is not entirely clear but recent studies raise the possibility that systemic OT may impact BAT thermogenesis through a direct mechanism. Yuan and colleagues found that slow continuous systemic infusion of OT (100 nmol/day) elevated rectal temperature (single time point), UCP-1 protein and UCP-1 mRNA in IBAT and IWAT of high fat diet-fed mice [24] where OTR are expressed [24]. In contrast, we found that a single acute bolus injection of lower doses of OT (5 and 10 μg/μL) elicited an initial reduction of T_IBAT_ prior to a subsequent elevation of T_IBAT_. Similarly, others have found that peripheral administration of higher doses of OT (1 mg/kg) elicited a robust hypothermic response [82]. Similar doses (1, 3 and 10 mg/kg, IP) were also found to block stress-induced stimulation of core temperature [83]. Kohli reported that pretreatment with arginine vasopressin receptor 1A (AVPR1A) antagonist can reduce OT-mediated hypothermia, while pretreatment with OTR antagonist does not [84]. Similar to systemic OT, we and others have demonstrated that systemic MTII produces an initial reduction of T_IBAT_ and/or core temperature [85; 86] followed by a subsequent elevation of T_IBAT_ or core temperature [86]. Like OT, MTII-elicited reduction of body temperature was found to be blunted in response to the AVPR1A antagonist [85]. In addition to a mechanism involving AVPV1A, MTII-elicited hypothermia is also due, in part, to mast cell activation [87]. Together, these findings suggest that the hypothermia in response to high doses of systemic OT might be mediated, in part, by arginine vasopressin (AVP) receptor V1A rather than through OTR. It remains to be determined whether the hypothermic effects of systemic OT are also mediated, in part, by activation of mast cells.

To our knowledge, this is the first time that SNS innervation of IBAT (NE content) has been found to be reduced in DIO rodents relative to age-matched lean control mice. These findings are consistent with the reduction of IBAT NE content in obese *fa/fa* Zucker rats relative to lean homo-(*Fa/Fa*) or heterozygous (*Fa/fa*) rats [88]. Similarly, tyrosine hydroxylase (TH; rate limiting enzyme to synthesis of catecholamines) was found to be reduced in obese relative to lean animals [89]. The same group also found a reduction of IBAT NE content, NE turnover and activity of dopamine-beta-hydroxylase (rate limiting enzyme to synthesis of NE) in young *fa/fa* Zucker rats prior to obesity onset [90]. Previous studies have found that the IBAT from humans with obesity appears to be hypoactive [91; 92] although it isn’t clear if this is due to reduced SNS innervation of IBAT or reduced sensitivity to endogenous catecholamines [93]. Consistent with these findings, obese offspring also show reductions of IBAT temperature [94]. Defects in BAT activity are associated with impairments of both the structure and function of BAT mitochondria [95]. Furthermore, the IBAT of obese animals was associated with enlarged lipid droplets [96], indicative of hypoactivity of IBAT. Together, our data support the hypothesis that impaired activation of IBAT in the context of DIO may be due, in part, to reduced SNS innervation of IBAT.

Our findings showing a reduction of IBAT UCP-1, Dio2, and Adbr3 in denervated mice is consistent with what has been reported from IBAT of denervated mice or hamsters [51; 52; 53; 54; 55]. In addition, we found a reduction of IBAT Adrb1, Acox1, UCP-3, bmp8b, Gpr120, and Cox8b in denervated mice. We also found that 4V OT blocked the reduction of Adrb1, Adrb3 and Dio2 in IBAT of denervated mice. Whether these genes contributed to the ability of OT to increase T_IBAT_ in denervated mice is unclear. What was also unexpected was that the beta-3 receptor agonist, CL 316243, was able to produce comparable effects on T_IBAT_ in both groups of mice despite there being a reduction of IBAT Adrb3. However, we have found that chronic 1x daily administration of CL 316243 produced similar effects on stimulation of T_IBAT_ on treatment day 1 vs day 19 despite there being a reduction of IBAT Adrb3 mRNA at the end of the study [97]. It is possible that the presence of even a reduced level of the beta-3 receptor in IBAT is sufficient to contribute to the effects of CL 316243 on IBAT temperature at the pharmacological doses used in our studies. It is not as likely that the effects of CL 316243 on IBAT temperature in denervated mice was due to action at other adrenergic receptors in IBAT because 1) CL 316243 is highly selective to the beta-3 receptor (128-fold higher selectivity to human beta-3 receptors vs beta-1 receptors [98]) and 2) is ineffective at increasing lipolysis, energy expenditure, and reducing food intake in beta-3 receptor deficient mice [99].

One limitation of our study is that restraint stress may have limited our ability to observe larger effects on T_IBAT_ [100] during the time period when the effect of the drug is relatively short-lived or small. We aimed to minimize the impact of restraint stress by adapting the animals to handling and mock injections during the week prior to the experiment and by administering the drugs during the early part of the light cycle when catecholamine levels [36] and IBAT temperature are lower [22]. Despite these adjustments to protocol, we failed to observe an obvious impairment in the ability of the sympathomimetic, tyramine, to stimulate T_IBAT_ in denervated mice even though there was clear evidence that IBAT NE content was lower in mice whose IBAT was denervated. While the effects of OT and CL 316243 continued well beyond the short-lived effects of restraint/vehicle stress on T_IBAT_ in our mouse studies (∼30-45 minutes), the effects of tyramine were relatively short-lived and may have been masked by restraint stress. Thus, stress-induced epinephrine, from the adrenal medulla, or global release of NE from SNS nerve terminals (in response to tyramine) [101; 102; 103], may have activated beta-3 receptors in IBAT to stimulate T_IBAT_, even in denervated mice.

We also acknowledge that the focus of this study was on IBAT given that IBAT is most well characterized BAT depot “because of its size, accessibility and clear innervation IBAT” [104]. However, IBAT is thought to contribute to approximately 45% of total thermogenic capacity of BAT [54] or ≥ 70% of total BAT mass [105]. Thus, it is possible that the other BAT depots (axillary, cervical, mediastinal and perirenal depots), all of which show elevated UCP-1 in response to cold [106], might have contributed to the effects of OT to elicit weight loss in IBAT denervated mice. Moreover, we acknowledge the potential contribution of IWAT and EWAT given that chronic 4V OT was able to elevate IWAT UCP-1 and EWAT Acox1 in a limited number of IBAT denervated mice. As mentioned earlier, previous findings demonstrate crosstalk between SNS circuits that innervate IBAT and WAT [51]. In addition, there is increased NE turnover and IWAT UCP-1 mRNA expression in IBAT denervated hamsters [51]. It will be important to 1) confirm our IWAT and WAT gene expression findings in a larger group of animals and 2) develop a model to assess the effectiveness of denervation of all BAT and specific WAT depots in order to more fully understand the importance of BAT and WAT depots in contributing to the effects of OT to elicit weight loss in rodent models.

In summary, our findings demonstrate that acute 4V OT (5 µg) produced comparable increases in T_IBAT_ in both denervated and sham mice. We subsequently found that chronic 4V OT produced similar reductions of body weight and adiposity in both sham and denervated mice. Importantly, our findings suggest that there is no change or obvious functional impairment in the response of the beta-3 receptor agonist, CL 316243, to activate IBAT in mice with impaired SNS innervation of IBAT in comparison to sham-operated mice. Together, these findings support the hypothesis that sympathetic innervation of IBAT is not a predominant mediator of OT-elicited increases in non-shivering BAT thermogenesis and reductions of body weight and adiposity in male DIO mice.

## ACKNOWLEDGMENTS

The authors thank the technical support of Nishi Ivanov. In addition, the authors are appreciative of the efforts by Dr. Michael Schwartz and Dr. Dianne Lattemann for providing feedback throughout the course of these studies.

Disclosures: JEB had a financial interest in OXT Therapeutics, Inc., a company developing highly specific and stable analogs of oxytocin to treat obesity and metabolic disease. The authors’ interests were reviewed and are managed by their local institutions in accordance with their conflict of interest policies. The other authors have nothing to report.

## Supplemental Study 1: Determine if surgical denervation of IBAT changes the ability of IP tyramine to increase T_IBAT_ in DIO mice

The goal of this study was to determine if IP tyramine-elicited increase in T_IBAT_ requires intact SNS outflow to IBAT in DIO mice. We selected doses of tyramine based on previous studies [102]. By design, mice were DIO as determined by both body weight (49.5±1.1 g) and adiposity (13.9±0.8 g fat mass; 31.2±1.4% adiposity) after maintenance on the HFD (60% kcal from fat; N=9-10/group) for approximately 4.25 months prior to sham/denervation procedures and implantation of temperature transponders underneath IBAT. Mice from **Study 4** were used in this study and were otherwise treated identically to those used in **Study 4**.

### Supplemental Study 1

In sham mice, tyramine (6.4 mg/kg) increased T_IBAT_ at 0.25-h post-injection while the higher dose (19.2 mg/kg) increased T_IBAT_ at 0.25 and 0.5-h post-injection (*P*<0.05; **Supplemental Figure 1A**). The lower dose also reduced T_IBAT_ at 0.75, 1, 1.25, 1.75, and 3-h post-injection and tended to reduce T_IBAT_ at 2-h post-injection. The higher dose reduced T_IBAT_ at 1, 1.25 and 1.75-h post-injection. Similar findings were apparent when measuring change in T_IBAT_ relative to baseline T_IBAT_ (**Supplemental Figure 1B**).

In denervated mice, tyramine (6.4 mg/kg) was not effective at increasing T_IBAT_ but tended to reduce T_IBAT_ at 0.75-h post-injection (*P*<0.05; **Supplemental Figure 1C**). The higher dose (19.2 mg/kg) stimulated T_IBAT_ at 0.25 and 0.5-h post-injection and reduced T_IBAT_ at 1.25, 1.5, 1.75 and 2-h post-injection (*P*<0.05). Similar findings were also apparent when measuring change in T_IBAT_ relative to baseline T_IBAT_ (**Supplemental Figure 1D**).

## Supplemental Study 2: Determine if surgical denervation of IBAT changes the ability of 4V OT to increase T_IBAT_ in lean mice

The goal of this study was to determine if OT-elicited increase in T_IBAT_ requires intact SNS outflow to IBAT in lean mice. By design, mice were lean as determined by both body weight (29.5±0.7 g) and adiposity (3.3±0.3 g fat mass; 10.4±0.8% adiposity) after maintenance on the chow (16% kcal from fat; N=10/group) for approximately 4-4.25 months prior to sham/denervation procedures and implantation of temperature transponders underneath IBAT. Mice were otherwise treated identically to those used in **Study 4**.

### Supplemental Study 2

Only a subset of samples from **Study 1B** were able to be screened for NE content but all mice were otherwise included in the analysis.

In sham mice, 4V OT (5 μg/μL) increased T_IBAT_ at 0.5, 0.75, 1 and 3-h post-injection (*P*<0.05; **Supplemental Figure 2A**) and tended to stimulate T_IBAT_ at 0.75 (1 μg/μL) and 1.25 (5 μg/μL) h-post-injection. In addition, we found similar findings were apparent when measuring change in T_IBAT_ relative to baseline T_IBAT_ (**Supplemental Figure 2B**).

Similarly, in denervated mice, 4V OT (5 μg/μL) increased T_IBAT_ at 0.75, 1, 1.25 and 3-h post-injection (*P*<0.05; **Supplemental Figure 2C**) and tended to stimulate T_IBAT_ at 1.25 (1 μg/μL) -h post-injection. In contrast, 4V OT was unable to stimulate a change in T_IBAT_ relative to baseline T_IBAT_ (**Supplemental Figure 2D**).

## Supplemental Study 3: Determine if a centrally (4V) effective dose of OT can increase T_IBAT_ when given into the periphery of lean mice

The goal of this study was to determine if systemic administration of OT (IP) can increase T_IBAT_ at a dose that was effective when given into the 4V in lean mice. By design, mice were lean (31.2±0.8 g) at study onset after maintenance on the chow (16% kcal from fat; N=10/group) for approximately 5 months. Mice did not undergo sham or SNS IBAT denervation procedures but were implanted with temperature transponders underneath IBAT and were otherwise treated similarly to those used in **Study 4**.

### Supplemental Study 3

In contrast to the more immediate elevation of T_IBAT_ in response to 4V OT (5 μg/μL), IP OT (5 μg/0.200 mL) resulted in a reduction of T_IBAT_ at 0.25-h post-injection followed by increases in T_IBAT_ at 1.25 and 3-h post-injection (*P*<0.05; **Supplemental Figure 3A**). IP OT (5 μg/0.200 mL) also tended to stimulate T_IBAT_ at 1.25, 2 and 4-h post-injection. In addition, we found similar findings were apparent when measuring change in T_IBAT_ relative to baseline T_IBAT_ (**Supplemental Figure 3B**).

In addition, a higher dose of OT (10 μg/0.200 mL) produced a similar reduction of T_IBAT_ at 15-min post-injection followed by increases in T_IBAT_ at 1.25, 1.5, 2, and 3-h post-injection (*P*<0.05; **Supplemental Figure 3A**). IP OT (10 μg/0.200 mL) also tended to stimulate T_IBAT_ at 105 min-post-injection. In addition, we found similar findings were apparent when measuring change in T_IBAT_ relative to baseline T_IBAT_ (**Supplemental Figure 3B**).

## Figure legend

**Supplemental Figure 1A-D: Effect of IP Tyramine on IBAT temperature (T_IBAT_) in DIO mice (0927.23F).** Mice were fed *ad libitum* and maintained on HFD (N=9-10/group) for 4.25 months prior to underdoing sham or SNS denervation procedures and implantation of temperature transponders underneath the left IBAT depot. Mice were allowed to recover for at least 1 week during which time they were adapted to a daily 4-h fast, handling and mock injections. Following completion of **Supplemental Study 5**, mice subsequently received IP injections (1.5 mL/kg; ≈ 0.07-0.09 mL/mouse) of tyramine (6.4 and 19.4 mg/kg) or vehicle (DMSO) where each animal received each treatment at least 48-h intervals. *A/C*, Effect of tyramine on T_IBAT_ in *A)* sham operated or *C)* IBAT denervated lean mice; *B/D*, Effect of tyramine on change in T_IBAT_ relative to baseline T_IBA_T (delta T_IBAT_) in *B)* sham operated or *D)* IBAT denervated DIO mice. Data are expressed as mean ± SEM. \**P*<0.05, †0.05<*P*<0.1 tyramine vs. vehicle.

**Supplemental Figure 2A-D: Effect of 4V OT on IBAT temperature (T_IBAT_) post-sham or IBAT denervation in lean mice.** Mice were maintained on chow (16% kcal from fat; N=10/group) for approximately for approximately 4-4.25 months prior to undergoing a sham or bilateral surgical IBAT denervation and implantation of temperature transponders underneath IBAT. Mice were subsequently implanted with 4V cannulas and allowed to recover for 3-4 weeks prior to receiving 4V injections of OT (1 or 5 μg/μL) or vehicle (sterile water) where each animal received each treatment at approximately 48-h intervals. *A/C*, Effect of OT on T_IBAT_ in *A)* sham operated or *C)* IBAT denervated lean mice; *B/D*, Effect of OT on change in T_IBAT_ relative to baseline T_IBA_T (delta T_IBAT_) in *B)* sham operated or *D)* IBAT denervated lean mice. Data are expressed as mean ± SEM. \**P*<0.05, †0.05<*P*<0.1 OT vs. vehicle.

**Supplemental Figure 3A-D: Effect of IP OT on IBAT temperature (T_IBAT_) in lean mice.** Mice were maintained on chow (16% kcal from fat; N=9-10/group) for approximately for approximately 2.5 months prior to implantation of temperature transponders underneath IBAT. Mice were subsequently implanted with 4V cannulas and allowed to recover for approximately 2 weeks prior to receiving acute 4V injections of OT or vehicle (data previously published [22]). Mice subsequently received IP injections of OT (5 μg/0.200 μL or 10 μg/0.200 μL) or vehicle (sterile water) where each animal received each treatment at approximately 48-h intervals. *A/C*, Effect of *A)* lower and *C)* higher dose of IP OT on T_IBAT_ in lean mice; *B/D*, Effect of *B)* lower and *D)* higher dose of IP OT on change in T_IBAT_ relative to baseline T_IBA_T (delta T_IBAT_) in lean mice. Data are expressed as mean ± SEM. \**P*<0.05, †0.05<*P*<0.1 OT vs. vehicle.

