## Supplementary figures and images for "Sympathetic innervation of interscapular brown adipose tissue is not a predominant mediator of oxytocin-elicited reductions of body weight and adiposity in male diet-induced obese mice"

### Supplemental Figure 1

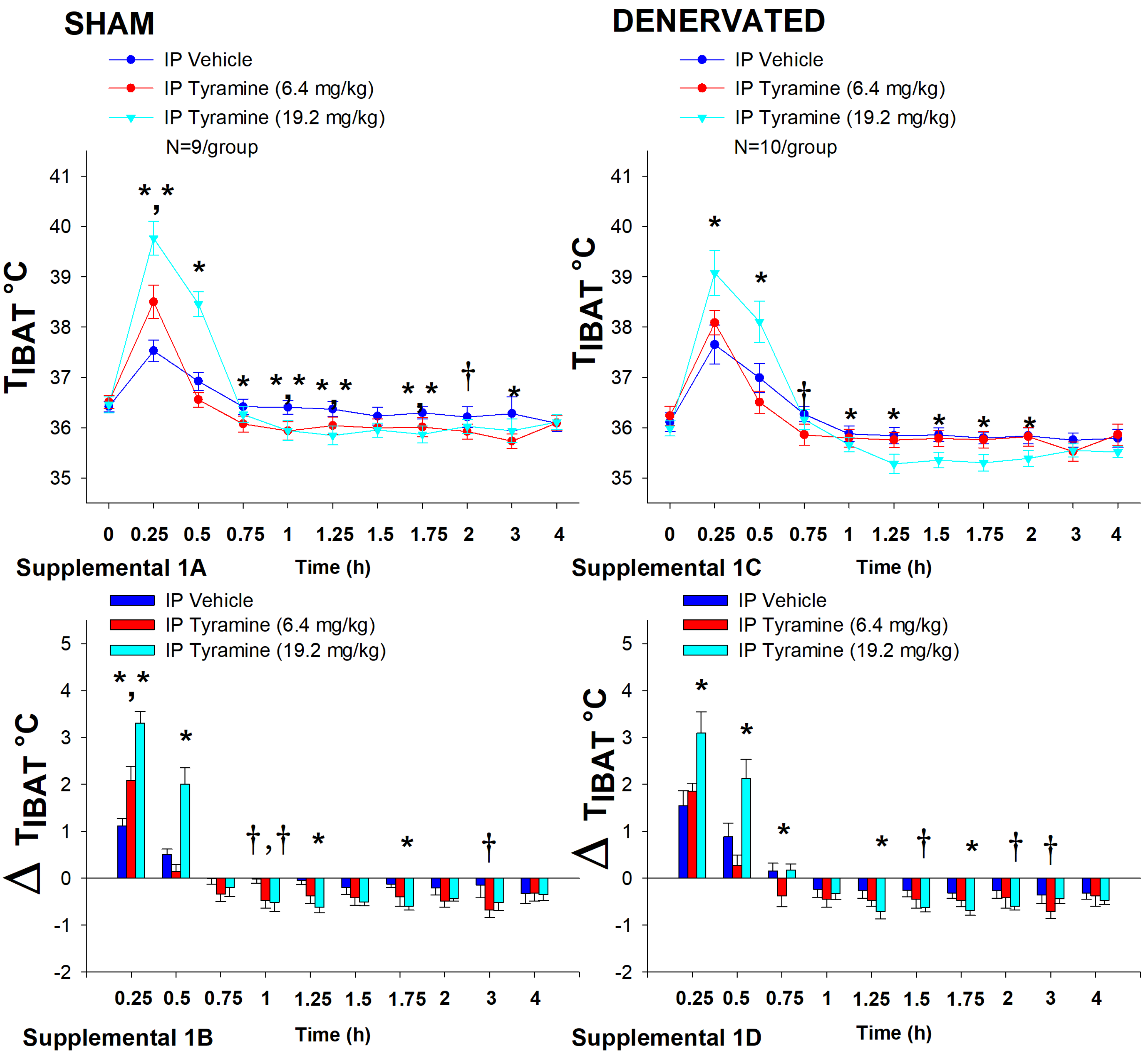

### Supplemental Figure 3

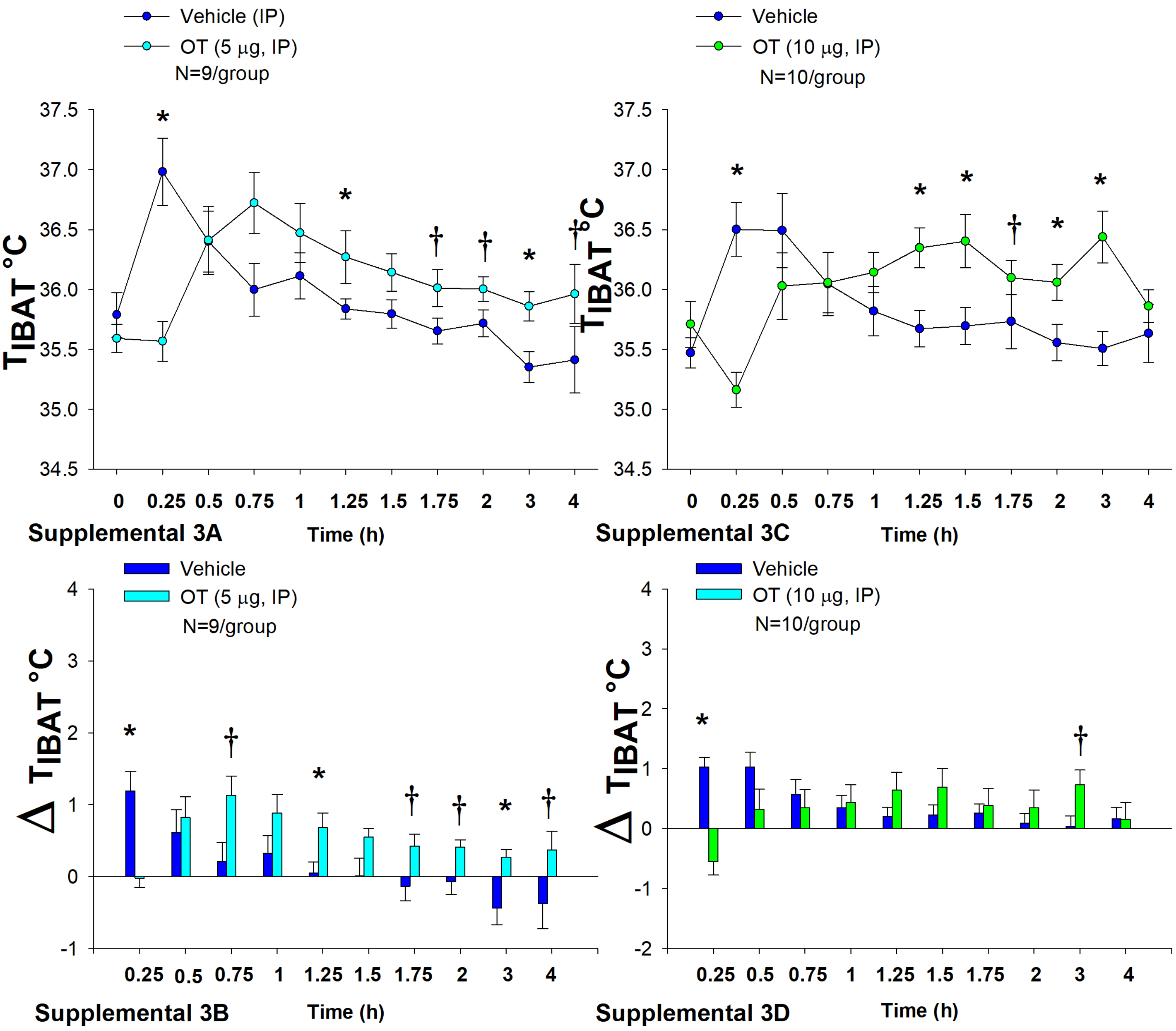
