## Supplemental Figure 2 for "Sympathetic innervation of interscapular brown adipose tissue is not a predominant mediator of oxytocin-elicited reductions of body weight and adiposity in male diet-induced obese mice"

### Effects of Acute 4V Oxytocin on IBAT Temperature in Lean Mice with Intact or Denervated SNS Outflow to IBAT

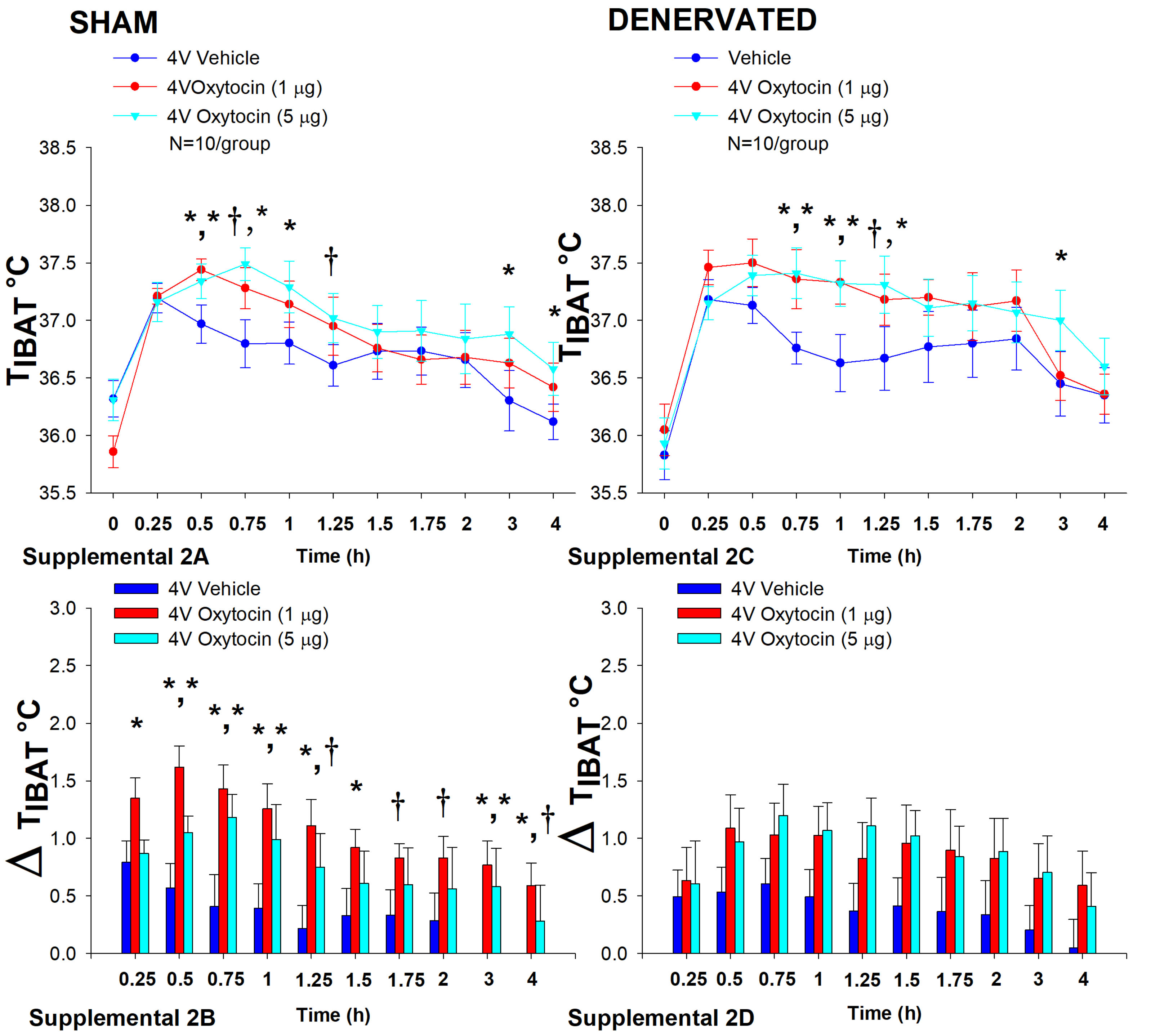
